# Selective profiling of translationally active tRNAs and their dynamics under stress

**DOI:** 10.64898/2026.03.02.709006

**Authors:** Mie Monti, Hasan Yilmaz, Alessia Del Piano, Michele Arnoldi, Isabelle Bonomo, Laia Llovera, Jessica Sarabando, Daniela Ribeiro, Ana Raquel Soares, Massimiliano Clamer, Eva Maria Novoa

**Affiliations:** Centre for Genomic Regulation (CRG), The Barcelona Institute of Science and Technology, Dr. Aiguader 88, Barcelona 08003, Spain; IMMAGINA Biotechnology, Trento, Italy; Institute of Biomedicine (iBiMED), Department of Medical Sciences, University of Aveiro, Aveiro, Portugal; Universitat Pompeu Fabra (UPF), Barcelona, Spain; 5ICREA, Pg. Lluís Companys 23, Barcelona, Spain

**Author notes:** Equal contribution (Mie Monti, Hasan Yilmaz). Correspondence to: Massimiliano Clamer and Eva Maria Novoa.

## Abstract

During translation, transfer RNAs (tRNAs) deliver specific amino acids to the ribosome in a coordinated manner with the sequence encoded by the mRNA. Despite their central role in protein synthesis, their precise contribution to the modulation of translation remains poorly understood, primarily due to lack of methods to characterise tRNA abundances and their modifications from actively translating ribosomes. Here we develop tRIBO-seq, a simple and robust nanopore-based method to selectively capture ribosome-associated native tRNA populations (ribo-tRNAs) from actively translating ribosomes, providing tRNA abundance, modification and fragmentation information from a single experiment. Using tRIBO-seq, we find that ribo-tRNAs, but often not total tRNAs, are significantly altered upon stress. Notably, we find that tRNAome alterations strongly vary depending on the stress type: while viral infection and leucine deprivation leads to changes in ribo-tRNA abundances, methionine starvation causes a dramatic loss of methyl-based tRNA modifications. By contrast, we find that arsenite exposure does not alter tRNA abundances nor modification patterns, but rather causes major fragmentation events in selected subsets of ribo-tRNAs. Altogether, tRIBO-seq offers a robust and reproducible approach to map the full tRNA landscape –capturing tRNA abundance, modification and fragmentation patterns–, across both total and actively translating tRNA populations, revealing the dynamics of the tRNAome with unprecedented resolution, in a single experiment.

## INTRODUCTION

The central dogma of molecular biology describes the flow of genetic information from DNA to messenger RNA (mRNA) and then to protein. Yet, in eukaryotes, the correlation between mRNA abundance and protein levels is often low^1,2^. This discrepancy highlights the importance of post-transcriptional regulatory mechanisms that influence protein synthesis beyond mRNA abundance alone^3^.

Transfer RNAs (tRNA) are molecular adaptors that translate genetic information encoded in mRNAs into the corresponding amino acid sequences in proteins ^4–6^. The anticodon of each tRNA molecule pairs with the corresponding codon of the mRNA, ensuring that the correct amino acids are delivered to the ribosome for incorporation into the growing polypeptide chain. To fulfill their function, tRNAs are decorated with a broad diversity of RNA modifications that are critical for stabilizing their structure, maintaining codon-anticodon interactions, and preventing frameshift mutations during translation ^7–9^. tRNAs are extensively modified, with an average of 13 modifications per tRNA molecule in eukaryotes ^10^. Different positions within the tRNA molecule can be subjected to modification, and different modification types can be present in the same tRNA molecule (e.g. acetylation, isomerization, methylation, thiolation) ^6,11^. This chemical diversity creates a vast combinatorial landscape for translational regulation, enabling cells to fine-tune translation in a context- and condition-dependent manner ^12–15^. Indeed, previous works have shown that certain tRNA modifications, such as methylation or thiolation, can be altered under specific stress exposures ^16–19^. Therefore, tRNAs are increasingly recognized as modulators of protein synthesis, fine-tuning the translational output ^10,18,20–24^.

To identify which tRNA modifications are dynamically regulated, highly sensitive assays such as liquid chromatography-mass spectrometry (LC-MS) ^19,25^ can be used to identify and quantify tRNA modification changes; however, these methods typically cannot identify which tRNAs and in which specific location modifications are altered. To overcome this limitation, targeted modification-specific sequencing techniques have been developed ^17,26–31^, but these are still blind to the majority of tRNA modifications, and can only profile a subset of tRNA modifications at a time –mainly those that alter the Watson-Crick base pairing moiety. More recently, advances in nanopore direct RNA sequencing (DRS) technologies have made it possible to capture native tRNA populations (Nano-tRNAseq), allowing for simultaneous assessment of tRNA abundance and modification patterns in a simple and high-throughput manner ^32^. Nano-tRNAseq relies on extending the tRNA molecules from both ends ^32–34^, thus retaining the information of the full-length tRNA molecule. Notably, nanopore-based tRNA sequencing methods circumvent the need for separate modification and abundance quantification assays, allowing for a more comprehensive analysis of tRNA dynamics under diverse conditions in a single experiment ^32,35,36^.

Previous works characterizing tRNA populations and their dynamics across conditions have typically focused on characterizing the total tRNA populations in the cell ^27,32,37–41^. However, total tRNA populations under nutrient-limiting or stressed conditions may not reflect the translational status of the cell, as they differ from ribosome-associated tRNA populations purified from polysomal fractions ^17,42,43^. While ribosome-associated-tRNAs are expected to more accurately reflect the translational status of cells, their isolation remains tedious and time-consuming, and requires large input amounts. Therefore, an efficient and simple method to capture actively translating ribosome-associated tRNAs is essential for understanding how the tRNAome is rapidly tuned to adapt proteomic output to environmental cues.

Here, we develop tRIBO-seq, a novel nanopore-based high-throughput method able to simultaneously capture tRNA abundances and modification information from both total (total-tRNAs) and ribosome-associated tRNA populations (ribo-tRNAs). Notably, we show that these two populations are largely identical under nutrient-rich, steady-state conditions. By contrast, we find that upon stress conditions, tRNAs are differentially recruited to ribosomes, revealing unique tRNAome adaptations that occur differently in ribo-tRNA and total-tRNA pools.

Using tRIBO-seq, we reveal that the tRNAome is differentially reprogrammed depending on the cellular stress: i) during viral infection, ribosome-associated tRNAs change in accordance with altered codon demand; (ii) upon arginine or leucine deprivation, we observe altered tRNA abundances of specific tRNA isoacceptors; iii) upon methionine deprivation, we identify global tRNA hypomethylation patterns; and iv) upon arsenite exposure, we observe selective tRNA fragmentation that mainly occurs in ribosome-embedded tRNAs, in agreement with recent works suggesting that angiogenin is activated at the ribosome ^44^. Notably, our work demonstrates that tRNA-derived fragments (tRFs) can be captured using nanopore-sequencing, providing a novel framework to study both tRFs and full-length tRNAs in a single platform and experimental sample. Altogether, our work demonstrates that tRIBO-seq is a simple and robust method, able to capture the tRNA pool and its dynamics from actively translating ribosomes, revealing changes in tRNA abundance, modification status, and fragmentation patterns upon diverse environmental exposures, from a single experiment.

## RESULTS

### Sequencing translationally-active tRNAs

tRNA dynamics has been predominantly examined using total tRNA (total-tRNA) populations, largely due to the burden and technical challenges posed by isolating ribosome-associated tRNAs (ribo-tRNAs), which typically requires labor-intensive sucrose gradient fractionation. However, total-tRNA and ribo-tRNA profiles can significantly diverge under various stress conditions, including as nutrient deprivation, chemical insults or viral infection ^42,43^. Therefore, ribo-tRNAs are expected to better represent the translational status of cells, however, their isolation remains cumbersome and time-consuming. Moreover, when ribo-tRNAs have been analyzed, the focus has primarily been on tRNA abundances^42,43^, leaving potential differences in post-transcriptional tRNA modification between total- and ribo-tRNAs largely unexplored.

To overcome this limitation, we developed tRIBO-seq, a novel method that can provide simultaneous quantitative measurements of both tRNA abundance and modification patterns both from tRNAs actively engaged in translation and total tRNA pools. To specifically sequence ‘active’ tRNAs, ribosomes were isolated using a puromycin derivative (RsP)^45^ followed by an antibody-free and tag-free pull-down (see *Methods*), thus obtaining the ribosome-bound fraction in a single step (**Fig. 1A**, see also **Fig. S1** and **Table S1**). Then, the small RNA fraction was isolated from each pool, and used as input for ligation with splint adaptors^32^, required for nanopore tRNA library preparation and sequencing. Notably, the mRNA populations found in ribo-embedded (Ribo-mRNAs) and total RNA fractions (Total-mRNA) can also be isolated and sequenced from the same samples (**Fig 1B**).

**Figure 1.**
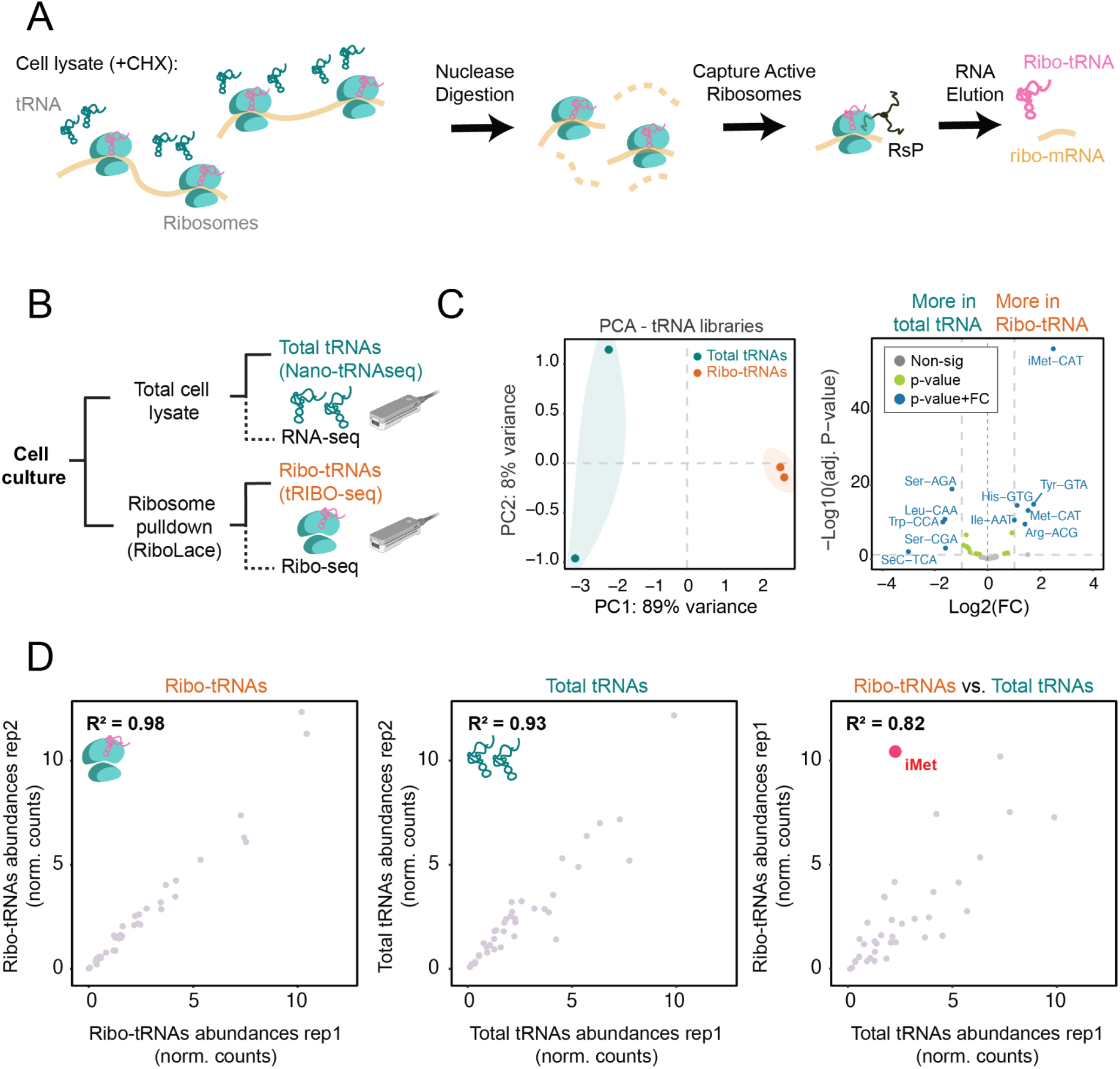
Experimental design and benchmarking of tRIBO-seq. **(A)** Schematic overview of the tRIBO-seq workflow. Translationally active ribosomes are isolated from cell lysates pre-treated with cycloheximide (CHX) to preserve ribosome-mRNA-tRNA complexes. Following nuclease digestion, ribosomes are purified using the RiboLace probe (‘RsP’) followed by RNA elution. The resulting RNA is used for obtaining ribo-tRNAs and mRNA footprints (Ribo-mRNAs), which can be sequenced using Ribo-seq. **(B)** Schematic representation of the experimental design: total tRNAs isolated from whole-cell lysates (total-tRNAs, captured using Nano-tRNAseq), and ribosome-associated tRNAs obtained from pulldown fractions (Ribo-tRNAs captured using tRIBO-seq). **(C)** Left: Principal component analysis (PCA) of tRNA abundances reveals clear separation between total and ribosome-associated tRNAs. Right: Volcano plot showing differential tRNA abundance when comparing ribo-tRNAs and total-tRNAs from HEK293T cell cultures (*n = 2* independent biological replicates per condition). Significance was defined as p < 0.05 with an absolute log2 fold-change ≥ 1. Full-length reads were used for pairwise analysis (see also **Table S2**). **(D)** Scatterplots depicting the replicability of tRNA abundances between independent biological replicates of HEK293T cultures, for ribo-tRNAs (left) and total-tRNAs (middle). Right: comparison between ribo-tRNA and total-tRNAs. The Pearson R2 correlation is also shown. See also **Fig. S1**.

To benchmark the method, we first applied tRIBO-seq to HEK293T cell cultures grown under standard-growth conditions (see *Methods*). Principal component analysis (PCA) of tRNA abundances revealed separate clustering of ribo-tRNAs and total-tRNAs in the first PC (**Fig. 1C**, see also **Table S2**), suggesting that even under standard-growth conditions, these populations are slightly distinct. To identify which specific tRNAs were responsible for the separation of the two populations, we performed differential expression (DE) analysis of tRNA abundances. We found that the initiator methionine tRNA (iMet-CAT) was the most significantly enriched tRNA in the ribosome-embedded fraction, consistent with its essential role in translation initiation^46^. Notably, tryptophan (Trp-CCA) and selenocysteine tRNA (SeC-TCA) appeared significantly decreased from the ribosome-associated pool, which aligns with the well-established scarcity of both amino acids in the human proteome ^47,48^.

The tRIBO-seq libraries showed high reproducibility across biological replicates and strong correlation between ribo-tRNA and total-tRNA samples (**Fig. 1D**), supporting the robustness of the approach. We then compared tRNA modification landscape of ribo-tRNAs and total-tRNAs, finding only minor differences in their modification patterns, suggesting that modifications of ribo-tRNAs largely resemble those of total-tRNAs under standard-growth conditions (**Fig. S3A**). Unexpectedly, our analysis also revealed differences in the CCA tail completeness between ribo-tRNAs and total tRNAs. While ribo-tRNAs displayed complete CCA tails for all tRNAs, which are required for aminoacylation, a partial loss of the CCA was observed in total tRNAs (**Fig. S3B**), consistent with previous works ^49^. Altogether, our results suggest that under normal elongation conditions, the total tRNA pool largely reflects the translationally active subset, both in terms of tRNA abundance and modification status, indicating that tRNA supply is well-matched to cellular translational demands.

To orthogonally validate tRIBO-seq, we then sequenced HEK293T tRNA populations extracted from polysome-enriched fractions (poly-seq), obtained using well-established sucrose gradient ultracentrifugation and fractionation methods ^42^ (see *Methods,* **Fig. S2A-C** and **Table S1**). Comparative analysis of tRNA counts generated by the two approaches revealed strong concordance (Pearson’s *r* = 0.89, *p* < 0.001; **Fig. S2D-F,** see also **Table S3**). Notably, initiator Met-tRNA (iMet-tRNA) was highly abundant in both datasets –but not in the total tRNA pool–, further supporting that tRIBO-seq accurately captures translationally active tRNAs (**Fig. S2F**). Importantly, tRIBO-seq recovered ∼18-fold more RNA relative to input (37% recovery), compared to poly-seq (2% recovery) (**Table S1**). Thus, relative to classical polysome profiling-based methods, tRIBO-seq enables the investigation of ribosome-associated tRNA populations using substantially lower input amounts and shorter experimental workflows (**Fig S2A**) while selectively capturing the translationally active tRNOme.

### tRIBO-seq captures translation initiation dynamics

To further validate that tRIBO-seq preferentially captures the ribosome-embedded tRNA fraction, we tested its performance in cell cultures treated with harringtonine (HAR), a well-characterized translational inhibitor that binds to the ribosomal A-site and arrests elongation immediately after translation initiation, resulting in the accumulation of ribosomes at the start codon ^50^ (**Fig. 2A** upper panel, see also **Fig. S4A**). To this end, we first performed ribosome profiling experiments to confirm that HAR exposure led to accumulation of ribosomes at ATG start codon, which was evidenced by a strong accumulation P-site reads at start codons (**Fig. 2A**, bottom panels). We then performed differential expression analysis of ribo-tRNA abundances between HAR-treated and control cells, finding that ribo-tRNAs showed significant enrichment of initiator tRNA (iMet-CAT) in HAR-treated cells (**Fig. 2B** left panel, see also **Fig. S4B-D** and **Table S4**), consistent with the drug-induced block of ribosomes at start codons. By contrast, total-tRNAs did not show a significant change in iMet tRNA abundance –or of any other tRNA– confirming the specificity of tRIBO-seq for capturing ribosome-embedded tRNAs (**Fig. 2B**, right panel, see also **Fig. S4D,E**).

**Figure 2:**
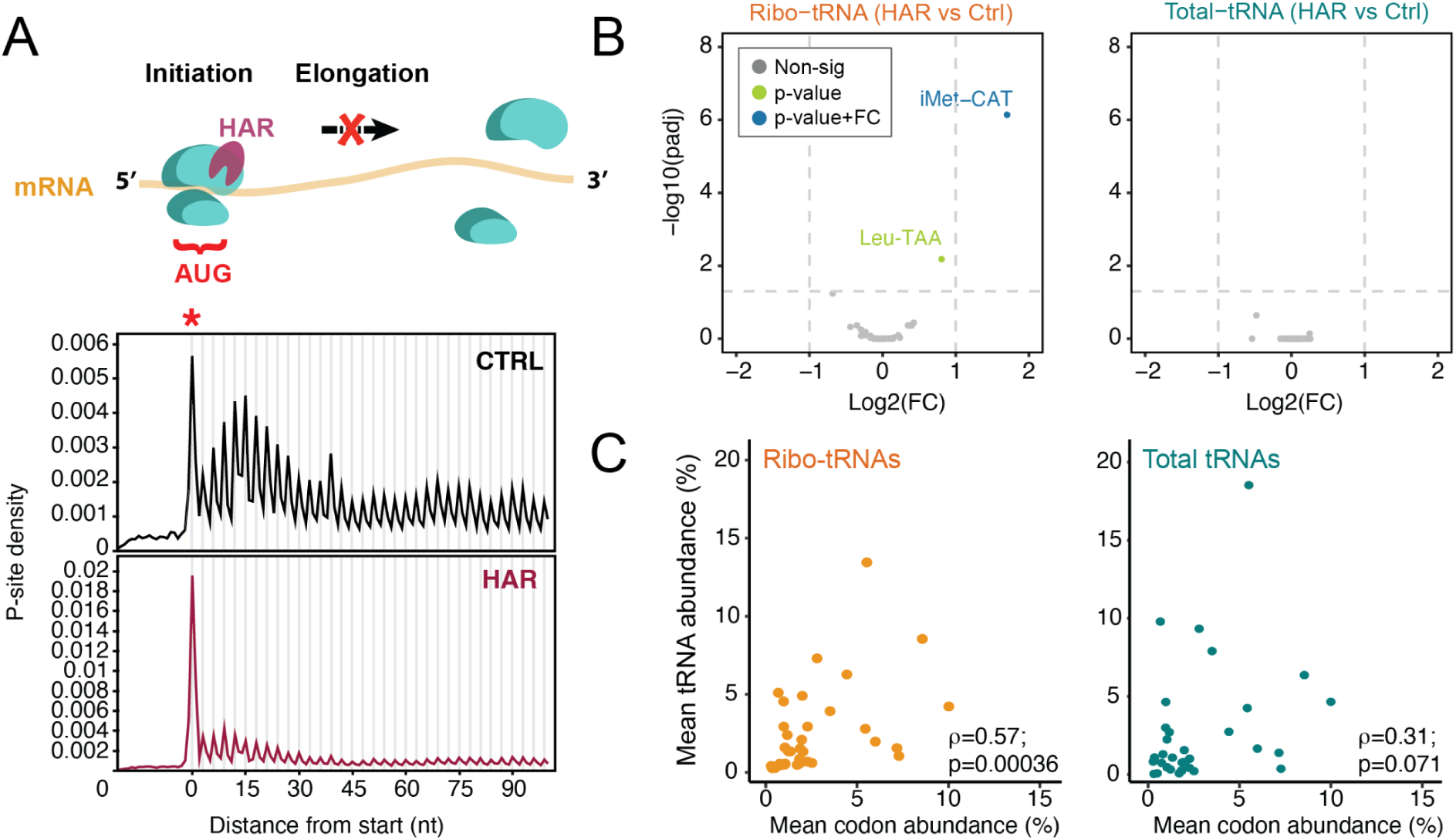
Validation of tRIBO-seq specificity using harringtonine. **(A)** Upper panel: Schematic representation of harringtonine (HAR) mechanism: HAR inhibits translation elongation, resulting in ribosome stalling at start codons. Bottom panel: Metagene plots obtained from ribosome profiling data in control (CTRL) and HAR-treated cells, showing an accumulation of P-site reads at start codons under HAR treatment. **(B)** Volcano plot depicting differential tRNA abundance between HAR-treated and control MCF-7 cells, for both ribo-tRNAs (left) and total tRNAs (right). Blue dots indicate tRNAs meeting both statistical significance and fold-change thresholds; green dots represent tRNAs meeting only statistical significance; gray dots correspond to non-significant changes. Significance was defined as p < 0.05 with an absolute log2 fold-change ≥ 1. Full-length reads were used for pairwise analysis. See also **Table S4**. **(C)** Scatterplots showing the correlation between mean codon abundance at the ribosomal P-site in control cells (x-axis) with tRNA abundances, using either ribo-tRNAs (left) or total tRNAs (right) populations (y-axis), respectively (n = 35). Spearman’s rho and associated p-values are also shown. All experiments were performed with *n* = 3 independent biological replicates per condition. See also **Fig. S4**.

Next, we examined the correlations between tRNA abundances of ribo-tRNAs and total-tRNAs with respect to the codon usage (see *Methods* and **Table S5**), which was quantified by extracting P-site codons from matched Ribo-Seq experiments on the same set of samples i.e., the control (elongating) condition. We found that ribo-tRNAs exhibited a significantly stronger correlation with codon usage at the ribosomal P-site compared to total-tRNAs, both when merging codons at the amino acid level (Spearman’s rho = 0.76 (ribo-tRNAs) and 0.57 (total-tRNAs) see also **Fig. S4F**), as well as using the Percudani rules^51^ for codon-anticodon matching (⍴ = 0.57 (ribo-tRNAs) and 0.31 (total-RNAs), see **Fig. 2C**). This enhanced codon-level concordance further validates the specificity of tRIBO-seq for capturing tRNAs that are functionally engaged with ribosomes during protein synthesis.

### Virus-driven codon demand is reflected in ribosome-associated tRNAs

We next asked whether tRIBO-seq could distinguish cellular responses to distinct forms of translational stress, such as viral infection or amino acid deprivation. During viral infection, viral mRNAs are expected to rapidly dominate ribosomal engagement, thereby reshaping cellular codon demand. To test whether tRIBO-seq captured viral-induced tRNA remodeling, we sequenced A549 cells infected with vesicular stomatitis virus (VSV), a negative sense RNA virus with a biased codon usage towards A/U-ending codons. RNA was collected at 6 hours post-infection, a time point characterized by active viral protein synthesis (**Fig. 3A**). Notably, previous studies have reported that polysome-associated tRNAs are significantly altered during infection with influenza A virus (a negative sense RNA virus), and during infection with vaccinia virus (a DNA virus), without detectable changes in total tRNA abundance ^42^.

**Figure 3:**
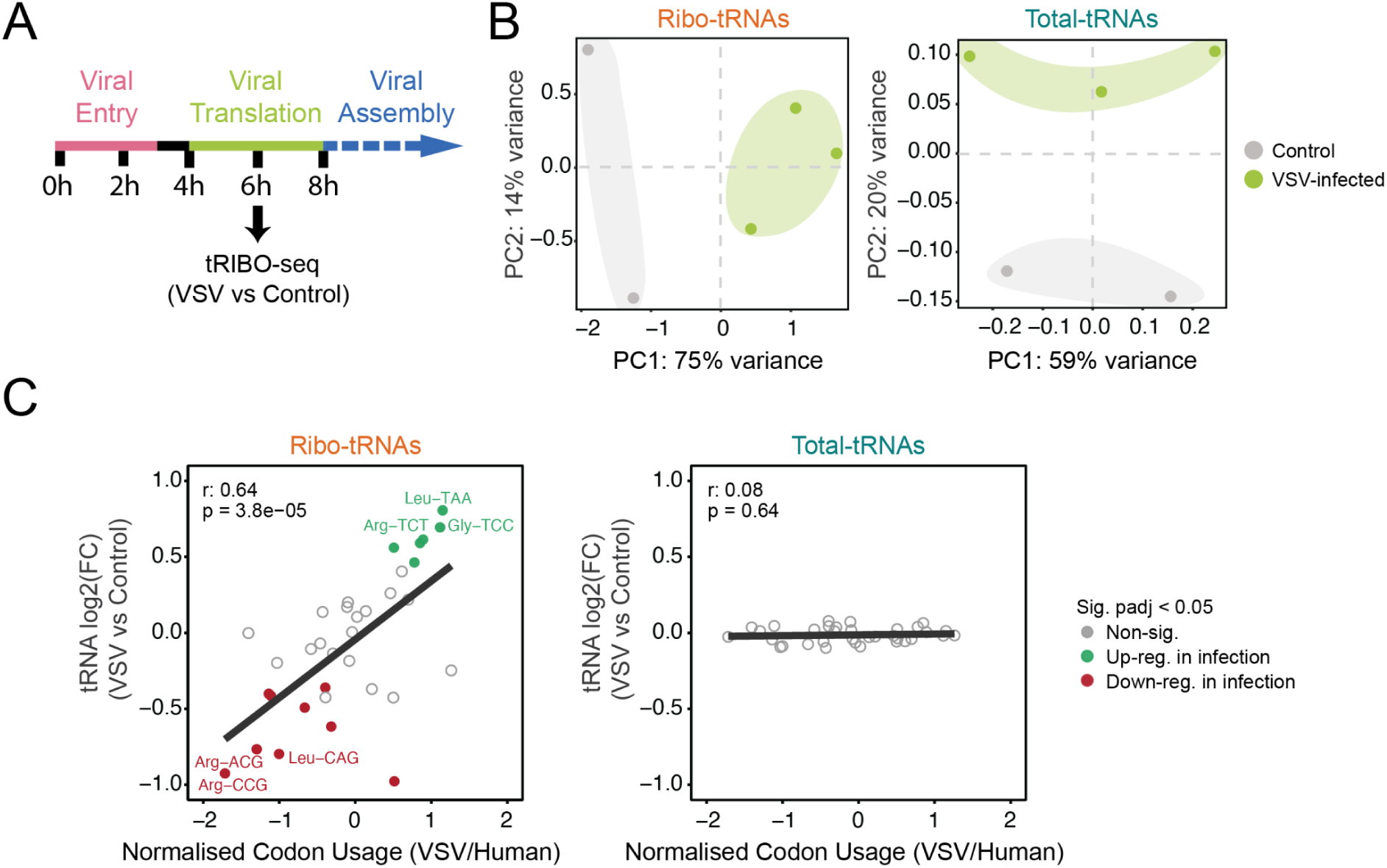
Virus-driven codon demand is reflected in ribosome-associated tRNAs. **(A)** Schematic timeline of vesicular stomatitis virus (VSV) infection. A549 cells were infected with VSV, and infected and non-infected control samples were harvested at 6 h post-infection, corresponding to active viral translation. Each condition was analyzed using three independent biological replicates, except for the non-infected control conditions (n = 2), due to insufficient sequencing coverage of one of the conditions. **(B)** Principal component analysis (PCA) of tRNA abundances for ribosome-associated tRNAs (ribo-tRNAs; left) and total tRNAs (right). Total tRNAs (right) are largely unaltered upon infection, which explains the poor clustering of the replicates based on their condition. By contrast, ribo-tRNAs (left) cluster separately along the PC1 based on their condition (VSV-infected vs control). **(C)** Scatter plots depicting the correlation between differential tRNA abundance (VSV vs Control) and VSV-to-human normalized codon usage. Ribo-tRNAs show a significant positive correlation, whereas no correlation is observed for total tRNA populations. Top three most up- and down-regulated tRNA genes labelled. Significance was defined as p < 0.05 with an absolute log2 fold-change ≥ 1. Full-length reads were used for pairwise analysis. Only tRNAs matched to their cognate codons according to Percudani rules were included (*n* = 38; see **Table S5**). See also **Table S6**.

Principal component analysis (PCA) of tRNA abundances revealed a clear separation between VSV-infected and uninfected control samples in the ribo-tRNA fraction along PC1, whereas total tRNA profiles obtained only separated on PC2 (**Fig. 3B**). This indicates that viral infection selectively alters the composition of ribosome-associated tRNAs without globally changing total tRNA levels.

To directly assess whether ribo-tRNA changes reflected viral codon usage, we correlated differential tRNA abundance with normalized VSV-to-human codon usage (see *Methods*). This analysis revealed a significant positive correlation for ribo-tRNAs (r = 0.64, p = 3.8 × 10⁻⁵), whereas no correlation was observed for total tRNA populations (**Fig. 3C**; see also **Table S6)**. These results show that tRIBO-seq specifically reports ribosome-associated tRNA redistribution in response to altered codon demand, and validate tRIBO-seq as a method to capture translationally engaged tRNAs.

### Ribo-tRNAs are sensitive to diverse amino acid deprivations

Previous works have shown that both arginine (Arg) and leucine (Leu) starvation can impair translation, albeit through distinct mechanisms –elongation stalling in the case of Arg starvation, and initiation inhibition in the case of Leu starvation ^52^ (**Fig. 4A**). To assess whether Arg or Leu deprivation would result in distinguishable ribo-tRNA profiles, we exposed HEK293T cells for 3 or 6 hours with media lacking either amino acid, using a fully supplemented medium as a matched control (**Fig. 4B**).

**Figure 4:**
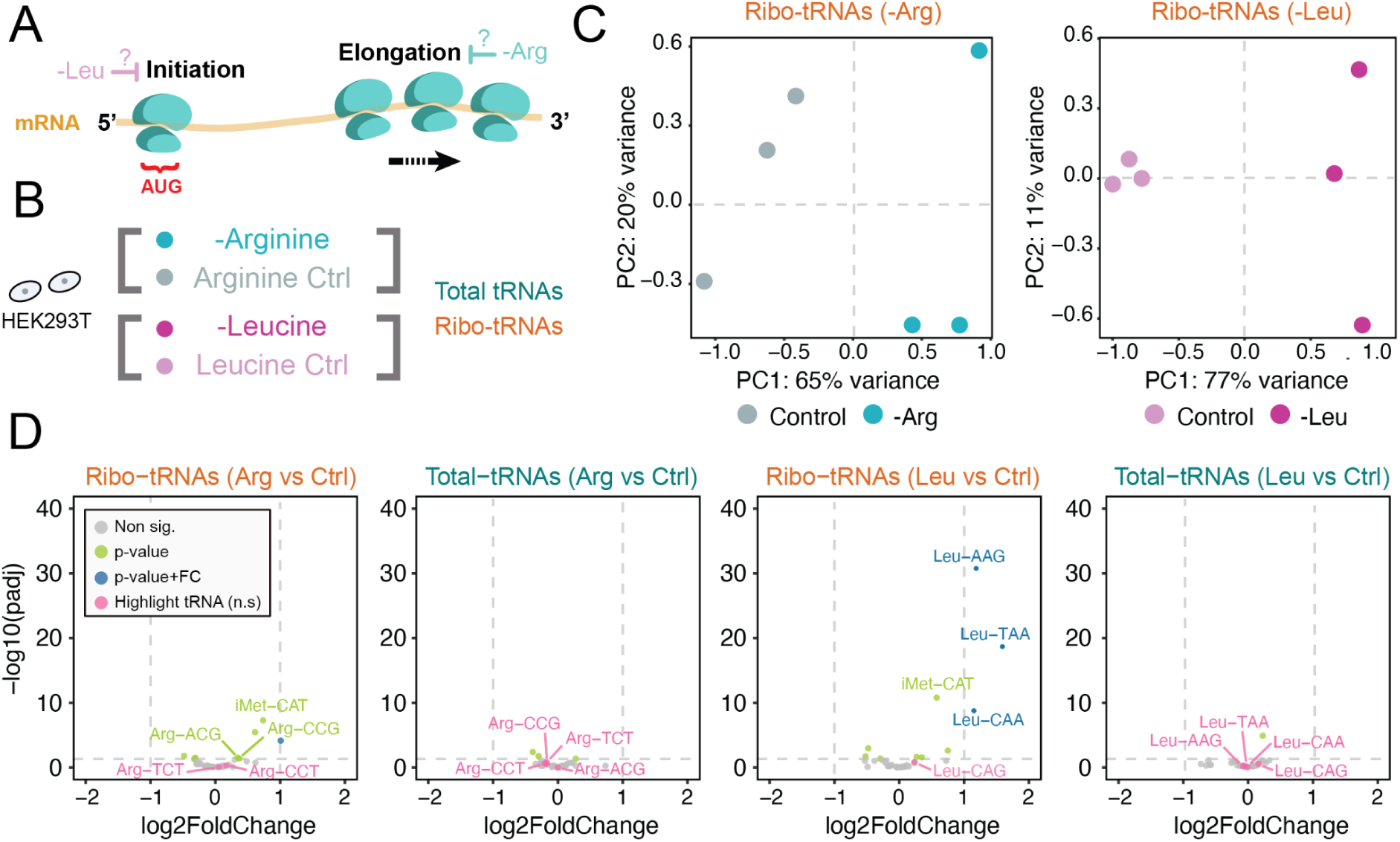
tRIBO-seq reveals distinct tRNA profiles under amino acid starvation. **(A)** Schematic of experimental rationale. Arg and Leu starvation perturb translation via different mechanisms: elongation stalling and initiation inhibition, respectively. **(B)** Schematic for the experimental design, in which HEK293T cells were cultured in amino acid deprived media for either Arg or Leu for 3 hours and their matching control media and subsequently analysed by both tRIBO-seq (ribo-tRNAs) and Nano-tRNA-seq (total-tRNAs). Each condition was performed with *n* = 3 independent biological replicates. **(C)** PCA of ribo-tRNAs shows separation of starved versus control samples along PC1, for both Arg (65% variance explained) and Leu (77%) starvation. **(D)** Volcano plots showing differential abundance of tRNAs in ribosome-associated and total pools for both starvation conditions. Significance was defined as p < 0.05 with an absolute log2 fold-change ≥ 1. Full-length reads were used for pairwise analysis. See also **Tables S8 and S9**.

Firstly, we confirmed that both Arg and Leu, with either 3-hour and 6-hour deprivation conditions, significantly affected the translational output using puromycin incorporation assays (**Fig. S5A**). This was further confirmed by polysome profiling experiments, which showed significant disruption of the polysome fraction following Arg and Leu deprivation, already at 3 hours (**Fig. S5B,C**). In addition, we performed ribosome profiling (Ribo-Seq) on Leu-deprived and control samples, which showed an increase in Leu codons at the A-site upon Leu starvation (**Fig. S6**; see also **Table S7**), suggesting that Leu starvation is already affecting ribosome elongation after 3-hour exposure.

Next, we sequenced ribo-tRNA and total-tRNA populations from 3h Arg- and Leu-deprived cells, as well as their control conditions (**Table S8 and S9**). PCA of ribo-tRNAs revealed a clear separation of starved versus control samples along the first principal component for both amino acid deprivation types (**Fig. 4C**). By contrast, total-tRNAs showed weaker clustering, indicating that ribo-tRNA composition strongly varies upon amino acid deprivation, whereas the global tRNA pool is only modestly affected by the amino acid deprivation (**Fig. S7A**). In both treated and untreated samples, we observed a strong enrichment in iMet-CAT tRNA when comparing ribo-tRNAs and total-tRNAs, both in Leu- and Arg-deprived cells (**Fig. S7B**), in agreement with our previous observations (**Fig. 1D and 2B**).

We then performed differential tRNA expression analysis of ribo-tRNAs from 3-hour Arg- and Leu-deprived compared to their control samples. This analysis revealed a significant enrichment of initiator methionine tRNA (iMet) in ribosomes from both Arg- and Leu-starved cells (**Fig. 4D**), in agreement with the increased monosome-to-polysome ratio detected in the polysome profiles (**Fig. S5B-C**), suggesting a translational shift toward initiation pausing or slowed elongation.

To our surprise, we also observed an enrichment of cognate tRNAs in the ribosome-associated pool under starvation, which contrasted with our initial expectation of depletion of these tRNAs in the ribo-tRNA pool upon amino acid starvation. This trend was most pronounced upon Leucine starvation, where several Leu tRNA isoacceptors were significantly enriched (**Fig. 4D**). In the Arginine condition, this pattern was more nuanced; only two isoacceptors, Arg-ACG and Arg-CCG, passed the significance threshold, but not the fold-change threshold. Extending the Arg deprivation up to 6 and 16 hours revealed a progressive increase of Arg-tRNAs significantly increased upon arginine starvation in the ribo-tRNA pool (**Fig. S8**, see also **Table S10**), supporting a slower but consistent response to arginine starvation. These results show that the counter-intuitive observation of Leu-tRNAs being significantly increased in ribo-tRNAs upon Leu starvation is likely extendable to other amino acids.

### Methionine starvation causes differential modification of ribo-embedded tRNAs

Given that amino acid deprivation can elicit distinct translational responses, we next asked whether methionine starvation induced changes in tRNA dynamics that resemble either leucine or arginine deprivation, or might instead elicit a distinct signature. Methionine (Met) is the initiator amino acid, and its deprivation has been previously shown to cause a drastic decrease in translation, assessed both by polysome profiling and puromycin incorporation assays ^53^. In addition, Met starvation has been shown to cause a strong transcriptional response ^54^. However, the effect of methionine starvation in the tRNA pool has so far not been explored. Here, we subjected MCF-7 cells to 6 and 16 hours of Met deprivation (**Fig. 5A**), and examined their tRNA profiles in both ribo-embedded and total tRNA populations, coupled with matched Ribo-seq and RNA-seq experiments.

**Figure 5.**
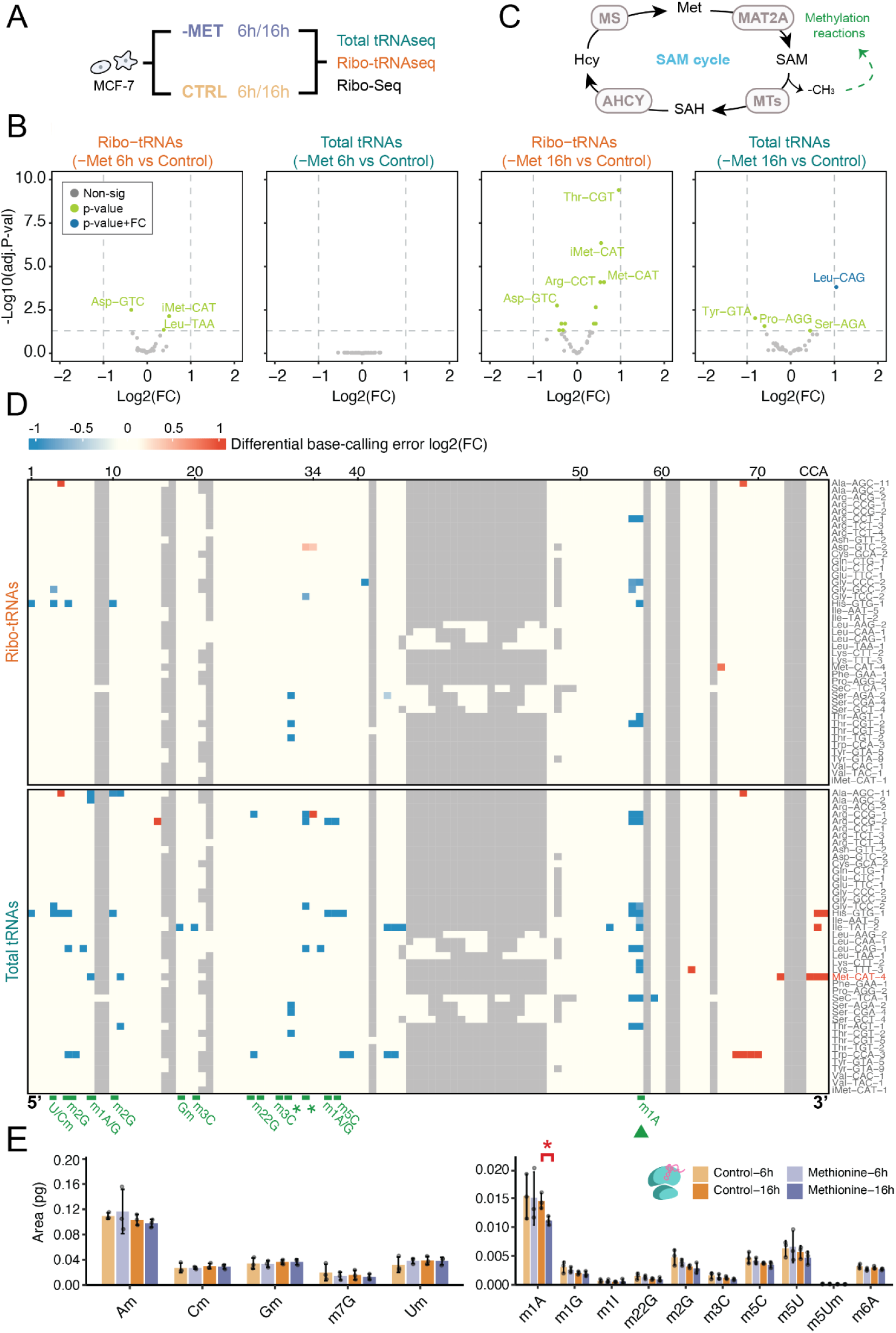
Methionine starvation causes a loss of tRNA methylation on tRNAs at 16h. **(A)** Schematic of the experimental design. MCF-7 cells were cultured in Met-deprived media for 6 or 16 hours, alongside matched controls, with *n* = 3 biological replicates per condition for all experiments. Total and ribosome-associated tRNAs were analyzed using tRIBO-seq (ribo-tRNAs) and Nano-tRNA-seq (total-tRNAs). **(B)** Volcano plots showing differential abundance of tRNAs in the ribosome-associated (ribo-tRNAs) and total pools (total-tRNAs), for both 6-hour and 16-hour Met starvation conditions. Significance was defined as p < 0.05 with an absolute log2 fold-change ≥ 1. For simplicity, only a subset of genes are labeled; complete results are provided in **Table S11**. **(C)** Schematic of the S-adenosylmethionine (SAM) biosynthesis and utilization cycle (see also **Fig. S11A**). **(D)** Heatmaps depicting differential modification (methionine-starved vs. control) for Ribo-tRNAs (top) and total tRNAs (bottom) at 16 hours. Differential basecalling errors are used as a proxy to quantify differential modifications across two conditions, as previously described^32,56^. The x-axis represents nucleotide positions (5’ to 3’); the y-axis lists tRNA isoacceptors (alphabetically ordered). Common methylation sites are annotated in green. See also **Fig. S12** for equivalent plots at 6h Met starvation. **(E)** LC–MS/MS quantification of modified nucleosides in ribo-tRNA isolated from methionine-starved and control cells. Areas were converted into picograms (pg) using standard curves for each nucleoside. Bars represent mean ± s.d. from *n* = 3 biological replicates per condition. P-values were calculated using a paired Student’s *t*-test comparing each condition to the corresponding control. Significance is indicated based on raw *p*-values. The left panel shows high-abundance nucleosides and the right panel low-abundance nucleosides. See also **Table S13**.

Firstly, we confirmed that Met deprivation yielded a strong transcriptional response both at 6 and 16h post-deprivation (**Fig. S9, Table S11**), in agreement with previous work ^54^. Despite the strong transcriptional response, analysis of the A-site codon occupancy from Ribo-seq data revealed no significant stalling at methionine codons (**Fig. S10E**), suggesting that most of the impairment occurs at the translation initiation stage, rather than at the elongation stage. In agreement with this, we found that the abundance of ribo-tRNAs and total tRNAs remained relatively stable upon Met starvation, with no significant changes at 6h post-deprivation and only modest changes in ribo-tRNAs at 16h (**Fig. 5B**, see also **Table S12**).

The set of modestly-upregulated ribo-tRNAs upon 16h methionine deprivation included methionine, arginine, glycine, serine, and threonine tRNAs (**Fig. 5B**). Notably, these amino acids are biochemically interconnected through the S-adenosylmethionine (SAM) biosynthesis pathway (**Fig. S11A**). SAM serves as a universal methyl donor for numerous cellular processes, including tRNA modification. In agreement with a previous report showing that methionine starvation enhances MAT2A expression via splicing regulation ^55^, we observed a robust upregulation of *MAT2A* upon methionine starvation (**Fig. S11B,C**). *MAT2A* is the enzyme catalyzing SAM synthesis from methionine (**Fig. 5C**). Notably, MCF-7 cells are deficient in MTAP ^54^, a key enzyme in the methionine salvage pathway, making them dependent on *de novo* SAM synthesis via *MAT2A* (**Fig. S11A**).

Given the observed dysregulation of SAM production pathways upon Met starvation, we wondered whether tRNA modifications requiring methyl groups would be significantly altered upon Met starvation. While no significant changes were observed at 6 hours (**Fig. S12B**), prolonged deprivation (16 hours) led to a strong and widespread hypomethylation of tRNAs, particularly in the total-tRNA pool, possibly reflecting reduced intracellular SAM availability. The hypomethylated positions overlapped with known methylation sites ^57^. By contrast, the effect in ribo-tRNAs was more modest (**Fig. 5D**), supporting the notion that proper tRNA modification is essential for their decoding function, as previously described ^21^. Finally, to further validate these observations, we performed nucleoside liquid chromatography-tandem mass spectrometry (LC-MS/MS) on control and Met-starved samples, finding that most methylation marks showed reduced trend in their modification levels, with m¹A being significantly decreased upon Met starvation at 16 hours (**Fig. 5E**).

### Oxidative stress triggers tRF formation in ribosome-associated tRNAs

We next asked whether an impairment of translation via chemical treatment would cause a distinct tRNA response than amino acid deprivation. To this end, MCF-7 cells were treated with 1 mM sodium arsenite (Ars), a well-characterized inducer of oxidative stress that acutely inhibits translation by triggering the integrated stress response (ISR) ^58,59^ (**Fig. 6A**, see also **Fig. S13A, Table S14** and *Methods*).

**Figure 6:**
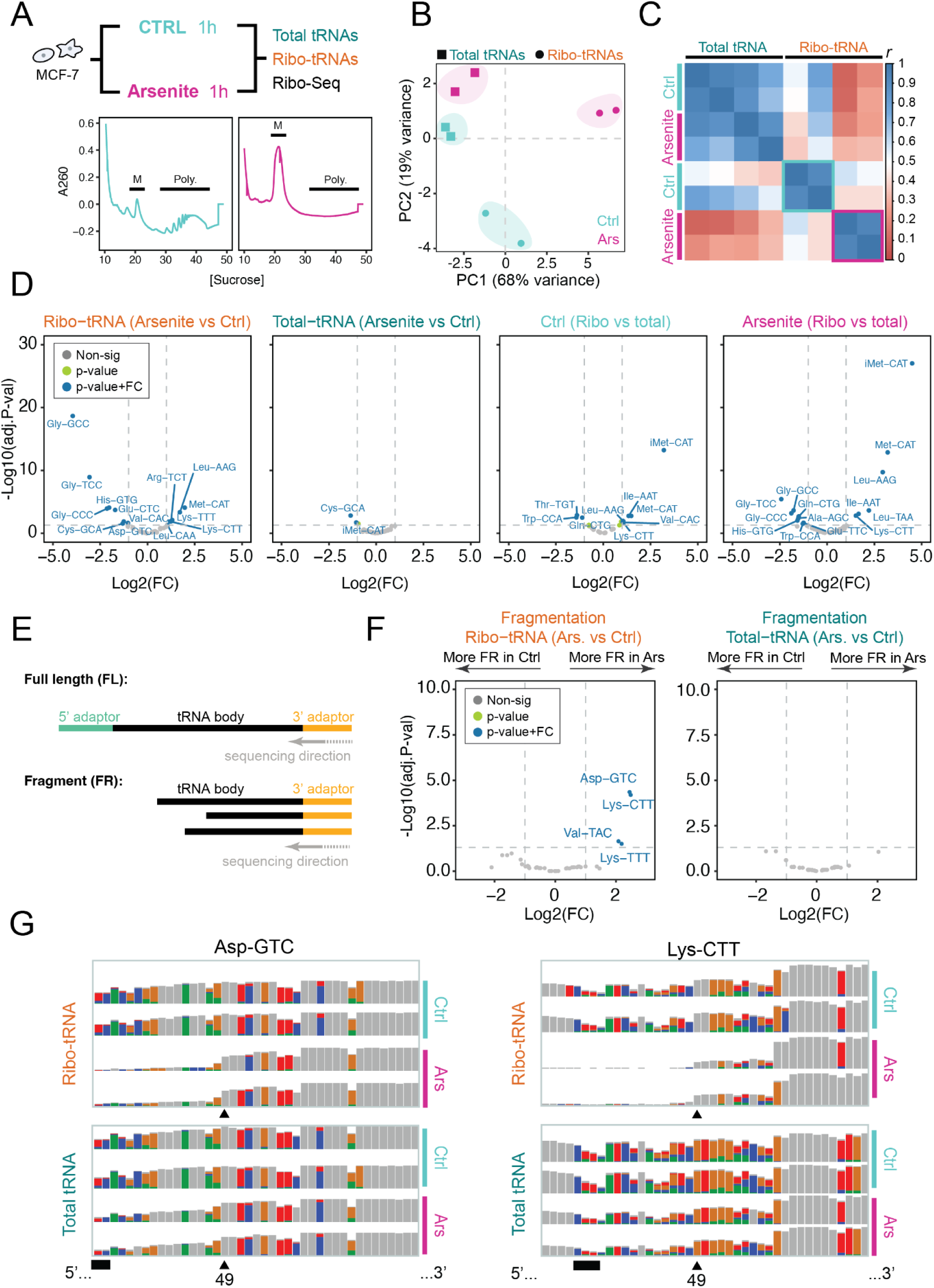
Selective fragmentation of Ribo-tRNAs in arsenite-treated samples. **(A)** Top: Schematic of the experimental design. MCF-7 cells were treated with 1 mM sodium arsenite for 50 minutes (Arsenite) or left untreated (CTRL). Total tRNAs and ribosome-associated tRNAs were profiled using Nano-tRNA-seq and tRIBO-seq, respectively. All experiments were performed with *n* = 2 independent biological replicates per condition. Bottom: Representative polysome profiles indicate translational repression following arsenite treatment. Peaks corresponding to monosomes (M) and polysomes (Poly.) are labeled. **(B)** PCA of total tRNAs and ribo-tRNAs. Arsenite treatment induces greater separation in ribo-tRNA profiles (circles) compared to total tRNA profiles (squares), indicating stronger effects on ribosome-associated tRNAs. **(C)** Pearson correlation matrix comparing tRNA abundance profiles across all conditions and library types (total vs ribo-tRNA). **(D)** Volcano plots showing differential abundance of tRNAs across all conditions and library types (total vs ribo-tRNA). See also **Fig. S15.** Significance was defined as p < 0.05 with an absolute log2 fold-change ≥ 1. No full-length filtering was applied in the pairwise analyses, as tRNA fragmentation is expected upon arsenite treatment (see **Table S15**). **(E)** Schematic illustrating classification of full-length (FL) vs fragmented reads. FL reads contain both 5′ and 3′ adaptors; fragmented reads contain only a 3′ adaptor. **(F)** Volcano plots showing differential fragmentation (FR vs FL) between arsenite and control samples (see *Methods*). **(G)** IGV genome browser snapshots for two representative tRNAs showing arsenite-induced fragmentation. For each condition and tRNA pool, two independent biological replicates are shown. Arrow indicates a significant fragmentation site. Anticodon underlined in black. See also **Fig. S18-S20.**

We first performed polysome profiling to confirm that arsenite exposure caused a global shutdown of active translation (**Fig. 6A**). This translational shutdown was accompanied by transcriptional upregulation of ISR effectors, such as *HSPA*, *ATF3*, and *PPP1R15A* (**Fig. S13B**), indicative of translational repression and stress adaptation. Indeed, 1h arsenite treatment resulted in 966 genes being exclusively translationally regulated, compared to 19 transcriptionally regulated genes and 13 that were affected at both levels (**Fig. S14A**), suggesting that post-transcriptional mechanisms dominate the cellular response to Ars-induced stress, in contrast with the predominantly transcriptional response observed upon Met deprivation (**Fig. S14B**).

To assess how oxidative stress influenced tRNA dynamics, we performed tRIBO-seq in cells treated with arsenite and untreated controls (**Fig. 6A**, see also **Table S15**). PCA and correlation analysis revealed that ribo-tRNA abundances –but not total tRNA abundances–were strongly affected by arsenite treatment (**Fig. 6B-C**). Differential expression analysis revealed that 14 of ribo-tRNAs were significantly altered upon arsenite treatment, in contrast to only 2 changing in the total-tRNA pool. This suggests that oxidative stress primarily affects actively translating tRNAs (**Fig. 6D**), consistent with the strong translational phenotype induced by arsenite (**Fig. S14A**). Among the few differentially expressed total tRNAs, the initiator methionine tRNA (iMet) was significantly affected (**Fig. 6D**, ‘Total-tRNA (Arsenite vs Ctrl)’), potentially reflecting its sequestration in stalled pre-initiation complexes, in agreement with in ISR activation ^59^. Within the ribo-tRNAs, we observed consistent depletion of tRNAs decoding cysteine, glutamate, and glycine, all of which are precursors of glutathione (GSH), a key antioxidant ^60^ (**Fig. 6D** ‘Ribo-tRNA (Arsenite vs Ctrl)’, see also **Fig. S15**). This pattern suggests a stress-induced metabolic shift, possibly reflecting impaired tRNA charging in response to oxidative stress. Unfortunately, arsenite treatment disrupted codon periodicity of ribosome footprints, limiting our ability to assess codon-level translation dynamics (**Fig. S16**).

We then examined the tRNA modification profiles, finding that, in contrast to methionine deprivation, arsenite stress did not significantly alter the tRNA modification landscape, neither in ribo-tRNAs nor total-tRNAs (**Fig. S17**). However, closer examination of the IGV tracks (**Fig. S18**) revealed another phenomena: a drastic increase in tRNA-derived fragments (tRFs) in ribo-tRNAs upon arsenite treatment, relative to the control condition.

Arsenite is well-documented to induce tRNA fragmentation ^61–63^. To determine whether tRIBO-seq was capturing tRNA-derived fragments, we performed fragmentation analysis, in which we compared full-length tRNA counts to tRNA fragment (tRF) counts, for each tRNA species (**Fig. 6F** and **Fig. S19A-B,** see also *Methods*). This analysis revealed four tRNAs significantly enriched in the ribosome-associated fraction following arsenite treatment: Asp-GTC, Lys-CTT, Val-TAC, and Lys-TTT (**Fig. 6F**). To exclude the possibility that the observed enrichment could result from nuclease digestion during tRIBO-seq preparation (**Fig. 1A**), we applied the same fragmentation analysis across all datasets presented in this study. No significant enrichment was detected when performing fragmentation analysis in other treatment conditions (**Fig. S20**), suggesting the observations are not an artifact of digestion, and are largely unique to arsenite treatment.

To further dissect the nature of ribosome-associated tRNA fragments, we next asked whether specific cleavage sites could be identified along the tRNA body. To this end, we performed position-resolved fragmentation analysis (**Fig. S19C**, see also *Methods*). This revealed prominent cleavage sites in several tRNAs around position 49 (**Fig. 6G** and **Fig. S19C**) suggesting a non-random, site-specific degradation pattern. We should note that due to the nature of nanopore direct RNA sequencing, if the 5’ adapter is not ligated –which would be the case in tRNA fragments–, the read will finish ∼13-15 bases before the true end of the RNA molecule ^64,65^, suggesting that the true cleavage site corresponds approximately to position ∼34-36 of these tRNAs, which corresponds to the anticodon region (**Fig. 6G**, see also **Fig. S18)**. In conclusion, tRIBO-seq captures arsenite-induced tRNA cleavage (leading to tRNA halves formation), which we find preferentially occurs in the ribo-embedded tRNA fraction **(Fig. 6F-G**, see also **Fig. S18-19**). This result is in agreement with previous works reporting that angiogenin activation occurs in the ribosome, and thus is likely preferentially degrading tRNAs in the ribosome ^44^.

## DISCUSSION

For decades, we have sought to understand how cells tune their translational output in response to external cues. However, a simple, information-rich and cost-effective approach to study the active tRNAome has been lacking. Consequently, previous works have largely focused on charting the dynamics of total tRNA pools, which, as we show here, can significantly differ from those tRNAs being actively used for translation.

Here, we present ribosome-embedded nanopore tRNA sequencing (tRIBO-seq), a novel and robust approach that enables multidimensional resolution of tRNA abundance, modifications and fragmentomics. By selectively profiling ribosome-associated tRNAs, this method provides a focused snapshot of the tRNA pool actively engaged in translation, substantially increasing the granularity of the data obtained, particularly under stress conditions. Importantly, mRNA populations can be recovered and sequenced from the same ribosome-embedded samples (Ribo-Seq), enabling integrated analysis of codon-anticodon interactions with minimal batch effects. The protocol can be completed within 5 hours and requires substantially lower input material than classical sucrose gradient-based polysome approaches (**Table S1**). Specifically, 3-5 µg of ribo-embedded RNA, which corresponds to ∼5 million cells depending on the translational status of the cell ^45^, is sufficient **(Fig. S1** and **S2)**.

We examined the ribo-tRNA and total-tRNA landscape under four different conditions. Firstly, we examined both tRNA pools under steady-state, nutrient-rich conditions, finding that the ribosome-associated tRNA population largely mimics the total tRNA pool (**Fig. 1**), indicating tight coordination between tRNA supply and the cellular translational needs. This aligns with the idea that, under steady-state conditions, cells maintain an optimized tRNA repertoire to support efficient translation. Subsequently, we examined how different stress-specific alterations reshaped this coordination. To this end, we first examined tRNA dynamics during viral infection, finding that VSV infection induced a pronounced alternation of ribo-tRNAs that closely reflects viral codon demand. By contrast, no significant changes in total-tRNAs were observed upon viral infection (**Fig. 3**).

Next, we examined how amino acid deprivation would affect the tRNA pool. While 3 hours of Leu starvation already produced significant changes in Leu-tRNAs in the ribo-tRNA pool, longer exposures were required to see similarly increased abundances of Arg-tRNAs within ribo-tRNAs (**Fig. 4** and **S8**). By contrast, methionine deprivation did not impact tRNA abundance, but rather caused widespread hypomethylation of the total tRNA pool at multiple well-characterized tRNA-modified sites (**Fig. 5**). The ribo-tRNA pool was also hypomodified, albeit to a lesser extent, suggesting a coordinated cellular response to preserve translational fidelity, in which modified tRNAs are preferentially loaded on ribosomes –possibly mediated by preferential aminoacylation of fully-modified tRNAs.

Finally, we observed that under oxidative stress conditions, such as arsenite exposure, there is a major rewiring of the translational landscape, characterized by strong changes in ribo-tRNA abundances as well as generation of stress-induced tRNA fragments, primarily at the ribosome (**Fig. 6**). Taken together, these observations highlight the interplay between the tRNA landscape and translation during cellular adaptation to stress, although direct impact on protein synthesis remains to be established. Overall, our results demonstrate that tRIBO-seq is a robust and comprehensive method to simultaneously quantify tRNA abundances, modification landscapes, and fragmentation patterns from a single experiment, using a single sequencing platform and from a single sample.

In recent years, it has been proposed that the tRNAome is being rewired upon external environmental conditions ^16,66,67^. In agreement with previous works ^42^, here we find that across cellular models and environmental conditions, the total tRNAome remains largely unchanged upon external stimuli. By contrast, our analyses reveal that ribo-tRNA populations are selectively remodeled in a stimulus-specific manner. Overall, this work highlights the importance of examining both ribo-embedded and total tRNAome population dynamics in order to comprehensively understand how the tRNAome is used in translation and rewired upon environmental cues. Notably, tRIBO-seq greatly simplifies the analysis of both ribo-tRNA and total-tRNA populations from a single sample.

Yet, several caveats and limitations remain when applying this technique. Firstly, the performance of tRIBO-seq will vary depending on the translational activity of the cell type.

For example, neuronal progenitors may require higher amounts of input material (e.g. 100-200 ug of total RNA), compared to cancerous cell lines such as HEK or MCF-7 (10-20 ug total RNA). Secondly, the protocol has only been tested in mammalian cell lines; it could possibly be extended to non-mammalian eukaryotic samples with some optimizations and variations in the protocol, but this has not been tested. Finally, while tRIBO-seq can capture both full-length tRNAs and tRNA-derived fragments (tRFs), we should note that it will only capture 3’tRFs –but not 5’tRFs–, due to its bias towards preferentially ligating CC(A)-ended transcripts (see **Fig. S18**). However, tRIBO-seq can be used to study tRF presence and formation, while also capturing full-length tRNA abundances and modification patterns.

Despite these challenges, tRIBO-seq offers a robust, high-resolution, and accessible method to unravel the dynamic roles of tRNA in translation under physiological and stress conditions. The workflow allows for sample processing to sequencing within a single day, without reliance on expensive or specialized instrumentation. Its unique combination of speed, depth, and multi-layered information that is obtained, offers an opportunity to dive into both canonical and non-canonical roles of tRNA, including ribosome engagement, modification dynamics, and fragmentation activities. This versatility enables a broad range of applications, including the dissection of tRNA-mediated regulatory mechanisms across diverse biological contexts, the evaluation of *in vitro* performance and *in vivo* efficiency of therapeutic tRNAs ^68,69^ and the assessment of drug incorporation into actively translating ribosomes. As we continue to explore tRNA’s regulatory and therapeutic potential, methods such as tRIBO-seq can be of great use to decode the intricate choreography of translation.

## MATERIALS AND METHODS

### Cell culture

HEK293T (ATCC), MCF-7 (Michigan Cancer Foundation-7, ATCC), and A549 (CCL-185, ATCC) cells were maintained at 37°C in a humidified incubator with 5% CO₂. HEK293T cells were maintained in Dulbecco’s Modified Eagle Medium (DMEM, Sigma-Aldrich, cat. no: 10566016) supplemented with 10% FBS (Thermo Fisher, cat. no: 10082147). MCF-7 cells were cultured in DMEM (Thermo Fisher, cat. no: 10566016) supplemented with 10% heat-inactivated FBS (Thermo Fisher, cat. no: 10082147), 1% (v/v) penicillin-streptomycin (Thermo Fisher, cat. no: 15140122), and 1 mM L-glutamine (Thermo Fisher, cat. no: 25030081). A549 cells were maintained in DMEM (Gibco) supplemented with 10% heat-inactivated FBS (Gibco) and 1% penicillin–streptomycin (Gibco). For routine passaging, cells were washed with 1X phosphate-buffered saline (PBS), detached using 0.25% trypsin-EDTA, and centrifuged at 300×g for 5 minutes. Cells were seeded at appropriate densities depending on the experimental requirements. All cell lines were routinely screened for *Mycoplasma* contamination using Eurofins Scientific services.

### Chemical treatments

MCF-7 cells were seeded at 1 × 10⁶ cells per 10-cm dish and treated with 1 mM sodium arsenite (Ars) for 50 minutes or 2 µg/mL Harringtonine (HAR) for 3 minutes, followed by 10 µg/mL cycloheximide (CHX) for 5 minutes. In all cases, DMSO was used as a vehicle control at an equivalent volume. We confirmed that the chosen arsenite treatment conditions did not compromise MCF-7 cell viability or morphology (**Fig. S21**).

### Cell viability assay using CellTiter-Glo

MCF7 cells were seeded at a density of 1 × 10⁴ cells per well in a 96-well plate. After 24 hours, cells were treated with sodium arsenite (Ars, 1 mM, 50 min) or harringtonine (HAR, 2 µg/mL, 3 min), followed by a brief cycloheximide (CHX) treatment (10 µg/mL, 5 min at 37°C) to mimic the conditions used in other assays. Vehicle controls (DMSO or H₂O) were included, with or without CHX. As a positive control for cell death, cells were treated with 5% Triton X-100 for 10 min.Cell viability was then assessed using the CellTiter-Glo Luminescent Cell Viability Assay (G7570, Promega) according to the manufacturer’s instructions. Briefly, after 30 minutes of kit equilibration at 25°C, an equal volume of reagent was added to each well, and plates were incubated at room temperature for 10 minutes to stabilize the luminescent signal. Luminescence was recorded using a Varioskan microplate reader (Thermo Fisher Scientific).

### Virus propagation and purification

Wild-type (wt) vesicular stomatitis virus (VSV) was propagated in cultured cells. Virus stock infectivity (plaque-forming units per milliliter, pfu/mL) was determined by standard plaque assay on Vero cells. For virus production, cells were infected with wt VSV at a multiplicity of infection (MOI) of 0.001. Virus adsorption was carried out for 1 h at 37 °C, with gentle agitation for 10 min at room temperature. Following adsorption, the viral inoculum was removed and replaced with DMEM (Thermo #61965059) supplemented with 1% FBS (Thermo #10270106) and 1% penicillin–streptomycin (Thermo #15070063), and cells were incubated for 18 h at 37 °C.

Culture supernatants were collected and clarified by centrifugation at 500g for 5 min at 4 °C to remove cellular debris. The clarified supernatant was transferred to ultracentrifuge tubes and centrifuged at 23,800 rpm for 90 min at 4 °C. Viral pellets were gently resuspended in NTE buffer (0.1 M NaCl, 1 mM EDTA, 0.1 M Tris-HCl, pH 7.4) and allowed to dissolve overnight on ice. For further purification, resuspended virus was layered onto a 10% sucrose cushion prepared in NTE buffer and ultracentrifuged at 40,500 rpm for 60 min at 4 °C. The final viral pellet was resuspended in NTE buffer and incubated overnight on ice in a cold room.

### Vesicular stomatitis virus infection in A549 cells

To perform infection with VSV, A549 (CCL-185, ATCC) cells were seeded at a density of 4×10^6^ cells in a 10-cm dish. Twenty-four hours later, cells were washed once with PBS and infected with VSV-wt (1,5×10^9^ pfu/mL) at a MOI of 3, prepared in serum-free medium (SFM) [DMEM supplemented without FBS and Pen-Strep]. Cells were incubated with the viral inoculum for 10 minutes at room temperature under gentle shaking, followed by a 45-minute incubation at 37°C in 5% CO_2_ to allow viral adsorption. After this period, SFM supplemented with 10% heat-inactivated FBS was added, and cells were further incubated for 6 hours at 37°C in 5% CO_2.._

### Arginine and Leucine starvation

HEK293T cells were initially maintained in standard DMEM (Sigma-Aldrich, cat. no: 10566016) supplemented with 10% fetal bovine serum (FBS) and incubated at 37 °C in a humidified atmosphere containing 5% CO₂. For deprivation, cells were washed with pre-warmed PBS and transferred to custom arginine-free or leucine-free media for 3 hours. The deprivation media were prepared by supplementing amino acid-free DMEM (Genaxxon Bioscience, cat. no: C4150) with 3.5 g/L glucose (19.4 mM) and the following amino acids (final concentrations, in mM): glycine (0.40), cysteine (0.20), glutamine (3.97), histidine (0.20), isoleucine (0.80), lysine (0.80), methionine (0.20), phenylalanine (0.40), serine (0.40), threonine (0.80), tryptophan (0.08), tyrosine (0.39), and valine (0.80). Arginine-free medium was supplemented with leucine (0.80 mM), and leucine-free medium was supplemented with arginine (0.40 mM). Control medium contained both L-leucine (0.80 mM) and L-arginine (0.40 mM) at equivalent concentrations. All media were pH-adjusted to 7.2–7.4, sterile-filtered, and supplemented with 10% dialyzed FBS (Gibco, cat. no: A3382001) and 1% (v/v) penicillin–streptomycin (Thermo Fisher, cat. no: 15140122). Media preparation and handling were performed under sterile conditions. Formulation details and corresponding supplier information are provided in **Table S16**. HEK293T cells were chosen for arginine and leucine starvation experiments because their translational responses to these precise amino acid deprivations have been characterized in previous works ^52^, finding that HEK293T cells exhibit distinct regulatory programs involving initiation inhibition and elongation stalling, depending on the amino acid that is deprived (Leu or Arg).

### Methionine starvation

Methionine starvation experiments were performed in MCF-7 cells, because this line is MTAP-deficient, and previous works have shown its metabolic sensitivity to methionine restriction ^54^. MCF-7 cells at ∼70% confluency were washed with pre-warmed PBS and incubated in methionine-free medium (MFM) for 6–16 hours. MFM was prepared using methionine-free DMEM (Thermo Fisher, cat. no: 21013024), supplemented with 10% dialyzed FBS (Gibco, cat. no: A3382001) and 1% (v/v) penicillin–streptomycin (Thermo Fisher, cat. no: 15140122). Control medium was generated by supplementing MFM with L-methionine to a final concentration of 0.2 mM (Thermo Fisher, cat. no: J6190418). All experimental and control conditions were prepared in parallel and maintained under identical incubation parameters.

### tRIBO-seq: ribo-embedded tRNA isolation

Ribosomes were isolated using the RiboLace kit (Immagina Biotechnology S.r.l., cat. no: GF001-12), according to the manufacturer’s instructions. The kit optimized for lysates corresponding to an input of 0.4-0.9 absorbance units (AU) at 260 nm (A260), which serves as a proxy for total ribosome content. Absorbance was measured using a Nanodrop spectrophotometer (Nucleic Acid function), with the lysis buffer used as blank. Briefly, MCF-7, HEK293T and A549 cells were seeded at 1×10⁶, 1.5×10⁶ and 4×10⁶ cells per 10-cm dish, respectively, and allowed to reach 70-80% confluency before treatment (see ‘Cell Culture and Treatment’ for dosage details). Prior to harvesting, cells were incubated with 10 ug/mL (CHX, SIGMA cat. no. 01810) for 5 minutes at 37°C. Cells were then washed twice with ice-cold PBS containing 20 ug/mL CHX and resuspended in 300 uL Lysis Buffer (LB; Immagina #1BT0031) supplemented with sodium deoxycholate 10% (v/v), DNase I (1 U/uL, Thermo Scientific catalog no. 89836) and RiboLock RNase Inhibitor (40 U/uL, Thermo Scientific catalog no. EO0381). We should note that cells were pre-treated with cycloheximide (CHX) to preserve ribosome-mRNA-tRNA interactions during isolation, which stabilizes translating ribosomes by binding the E-site ^48^. Absorbance of the cell lysate was measured at 260 nm using a Nanodrop ND-1000 UV-VIS Spectrophotometer, and 0.9AU was used for ribosome pull-down, adjusted to a final volume of 450 uL with Wash Buffer (WB; Immagina #IBT0071). To each sample, 0.9 uL of Nux Enhancer (NE; Immagina #IBT0081) and 4.5 uL diluted Nuclease solution (equivalent to 6.75U, dNux; Immagina # IBT0081) were added, and the samples were incubated at 25°C for 45 minutes to allow for digestion. Digestion was terminated by adding 1.5 uL SUPERaseIn (Invitrogen, catalog no. AM2696) and incubating on ice for 10 minutes. While the digestion occurred, ribosome pull-down beads were prepared by functionalizing RiboLace magnetic beads (RmB; Immagina #IBT0042) with the RiboLace smart probe (RsP; Immagina # IBT0011). The probe was allowed to bind to the beads for 1 hour at room temperature with shaking at 1400 RPM. After incubation, mPEG (Immagina #IBT0061) was added to block non-specific binding, and the beads were washed twice with WB. The digested cell lysate was added to the functionalized beads and incubated for 70 minutes on a rotating wheel (3-10 RPM) at 4°C. Following incubation, the beads were washed twice with 1 mL WB. RNA was then eluted from ribosomes using the following RNA Clean & Concentrator^TM^-5 kit (Zymo catalog. No. R1015 or R1016) according to manufacturer instructions, with a modification: 200 uL of Zymo RNA Binding Buffer was added directly to the beads, followed by thorough mixing. Recovered RNA was quantified at 260 nm using a Nanodrop ND-1000 UV-VIS Spectrophotometer, and RNA integrity and size were assessed via 15% TBE-UREA gels and/or Agilent 4200 TapeStation RNA HS ScreenTape Assay (cat. no: 5067-5579).

### tRIBO-seq: library preparation

For building ribo-embedded nano-tRNA-seq libraries, the protocol of Lucas et al.^32^ was followed, with minor adaptations specific to ribo-embedded tRNA sequencing. Briefly, since CHX treatment retains deacylated tRNAs at the peptidyl (P) site, as described by Schneider-Poetsch et al. (2010), the deacylation step was omitted for these samples. However, as harringtonine (HAR) blocks translation before peptide bond formation between P-site and aminoacyl (A) site tRNAs (Dmitriev et al., 2020), HAR-treated libraries were deacylated to enable adaptor ligation for sequencing. Nano-tRNAseq libraries from total RNA were prepared following the protocol by Lucas et al. ^32^. All libraries were sequenced using the Oxford Nanopore Direct RNA Sequencing (DRS) protocol with the DRS kit (ONT, SQK-RNA002).

### tRIBO-seq: data analysis

#### Basecalling and mapping

Basecalling was performed using Guppy (v 6.5.7), as described by Lucas et al. ^32^. Basecalled reads were aligned to the reference genome using minimap2 (v2.24) ^70^ with the parameters: -ax splice -k 7 -w 3 -n 1 -m 13 -s 30 -A 2 -B 1 -O 1,32 -E 1,0. Demultiplexing was conducted using SeqTagger ^71^. Reads were subsequently separated based on their alignment positions to annotated reference tRNAs.

#### Filtering

For downstream analyses, reads lacking the 3′ splint adapter were discarded, as those most likely correspond to mismapped reads. Full-length reads were defined by their start positions relative to the reference tRNAs. Reads containing full or partial 5′ splint adapters were kept for downstream analyses, and were labeled as “full-length tRNAs” and “fragmented tRNAs”, respectively. All reads were further filtered based on mapping quality (MapQ ≥ 10). All filtering steps were carried out using pysam (v0.21.0) within custom Python scripts, available in GitHub (see ‘Data Availability’ section below).

#### Differential tRNA abundance and Principal Component Analysis (PCA)

Differential abundance analysis was conducted using DESeq2 (v1.44.0) within a custom R script (see ‘Data Availability’). tRNAs with fewer than 10 counts across all samples were excluded. Size-factor normalization from the DESeq2 ^72^ package was applied prior to differential testing. Comparisons between groups (Control vs. Treatment) within each experimental condition (Ribo-embedded vs. Total) were modeled using a one-factor design matrix: ∼ Group. P-values were adjusted for multiple testing using the Benjamini-Hochberg procedure ^73^. This analysis was performed separately for each of the three datasets: “All”, “Full-Length”, and “Fragment”. PCA was conducted using the same normalized DESeq2 data structure, variance data generated by the varianceStabilizingTransformation function, and PCA plots were generated using the plotPCA function from the DESeq2 package.

#### Differential modification analysis

Differential modification analysis was based on the detection of basecalling errors, as previously described by Begik et al. ^74^. Only the “Full-Length” subsample was used in this analysis to minimize false positives due to positional coverage variability. Positions with fewer than 10 total counts across all samples were excluded. For each reference tRNA, positions matching the expected nucleotide at each position were labelled as “not-modified,” while mismatches, insertions, or deletions were classified as “modified” counts. This was implemented via a custom R script (see ‘Data Availability’). Size-factor normalization from the DESeq2 ^72^ package was applied prior to differential modification testing. Differential modification analysis was performed using DESeq2 (v1.44.0), applying a two-factor design: ∼ Condition * Type, where ‘Condition’ refers to control versus treated and ‘Type’ refers to modified versus not-modified. P-values were corrected using the Benjamini-Hochberg method. Log₂ fold changes were visualized using ComplexHeatmap ^75^ (v2.22.0). Final heatmaps were generated filtering out positions with adjusted P-values > 0.05.

#### Fragmentation analysis

Fragmentation was assessed for each read by normalizing its length relative to reference length, yielding a fragmentation ratio. *Modification Analysis*: Read counts per position were computed using the ‘mpileup’ module from ‘pysam*’* within a custom Python script. For each tRNA position, the number of correctly and incorrectly called bases was summed and aggregated into a structured dataframe. This dataframe was then used for differential analysis in *DESeq2*, employing a two-factor model with “Condition” (Non-Treated vs Treated) and “Type” (Correct Call vs Incorrect Call) as factors. Resulting p-values were adjusted using the Benjamini-Hochberg method. The analysis aimed to identify positions with significant differences between conditions, which were interpreted as potential RNA modification sites.

### Ribo-seq library preparation

Library preparation of eluted ribosome protected fragments (RPFs) was performed using the RiboLace 360 Gel Free kit (Immagina Biotechnology S.r.l, cat. no: 360TE-12). Briefly, 1.5 ug of RNA (see ‘RiboLace-based ribosome isolation and RNA elution’) was 5’ phosphorylated by incubation with Buffer L1 (Immagina #IBT0151), ATP (Immagina #IBT0171), and enzyme L1 (Immagina #IBT0161) in a final volume of 50 µL for 1 hour at 37°C. The reaction mixture was allowed to incubate for 1 hour at 37°C. The reaction was cleaned to retain small RNAs using the RNA Clean & Concentrator-5 kit (Zymo, cat. no. R1015 or R1016). Adaptors were ligated to 5′ phosphorylated RNA, followed by incubation with Buffer L2 (Immagina #IBT0181), GTP (Immagina #IBT0201), MnCl₂ (Immagina #IBT0211), L2 enzyme (Immagina #IBT0191), and Linker MC+ (Immagina #IBT0222) for 1 hour at 37°C, then purified with the RNA Clean & Concentrator-5 kit. Circularization was achieved by mixing adaptor-ligated RNA with Buffer L3 (Immagina #IBT0231), ATP (Immagina #IBT0172), PEG8000 (Immagina #IBT0251), and enzyme L3 (Immagina #IBT0241), then incubating for 2 hours at 25°C, followed by purification to retain small RNAs. Reverse transcription was performed by incubating circular RNA with dNTPs (Immagina #IBT0301) and Primer L4 (Immagina #IBT0262) at 70°C for 5 minutes, cooling on ice, and then adding Buffer L4 (Immagina #IBT0271), DTT (Immagina #IBT0291), and enzyme L4 (Immagina #IBT0281). The reaction was incubated at 50°C for 40 minutes, heat-inactivated at 70°C for 10 minutes, then treated with AR enzyme (Immagina #IBT0311) at 37°C for 1 hour, followed by 80°C for 20 minutes. PCR amplification was conducted on 20 µL cDNA with L5 enzyme (Immagina #IBT0321), primers F1 (Immagina #IBT0331) and R1 (Immagina #IBT0341), and nuclease-free water. The program included an initial denaturation at 98°C for 1 minute, followed by 8 cycles of 98°C for 30 seconds, 61°C for 30 seconds, and 72°C for 10 seconds, with a final hold at 4°C. The product was purified with Agencourt AMPure XP beads (Beckman Coulter, cat. no. A63881). A second PCR for library multiplexing incorporated unique dual indices (UDIs) using previous PCR product, L5 enzyme (Immagina #IBT0321), and LACEseq UDIs (10 µM). The program included an initial denaturation at 98°C for 1 minute, followed by 6 cycles of 98°C for 30 seconds, 60°C for 30 seconds, and 72°C for 10 seconds, with a final hold at 4°C. The product was purified using the NucleoSpin Gel and PCR Clean-Up kit. The final library was evaluated with an Agilent 2100 Bioanalyzer using the High Sensitivity DNA kit. Sequencing was performed using the NovaSeq platform, achieving a coverage of 150 million reads per sample.

### Ribo-seq data analysis

Sequenced reads were pre-processed to trim adapter sequences using Cutadapt (version 4.9). Unique Molecular Identifiers (UMIs) were then extracted using UMI-tools (version 1.1.6), and contaminant reads corresponding to rRNAs, tRNAs, and ncRNAs were filtered out by pre-mapping to an artificial ‘contaminant’ genome using Bowtie2 (version 2.5.4). The pre-processed reads were then aligned to the human transcriptome (ensembl 108) using STAR (version 2.7.10b), and the resulting BAM files were sorted and indexed using Samtools (version 1.21). Ribosome profiling analysis was performed using Martian (version 1.5.0), using the GENCODE human database release 42. The lengths of ribosome-protected fragments (RPFs) corresponding to monosomes were identified by determining those aligning to the correct reading frame. The primary monosome length was defined as the most frequent RPF length aligning to the correct frame,and a 3-nt frame interval was selected as the monosome length range. Subsequent analyses were restricted to reads within this defined range. Additional methodological details can be found in the Martian documentation: https://github.com/ImmaginaBiotechnology/Documents/blob/main/Martian.md

### Codon-anticodon analysis

Codon-anticodon analysis was performed using the Percudani rules^51^ which assume that adenosine at the first (5′) position of the tRNA anticodon undergoes A-to-I (inosine) editing, enabling wobble pairing. In cases where adenosine is absent at this position, guanosine is assumed to wobble instead (see **Table S5**).

For codon usage analyses in viral experiments, codon usage tables for vesicular stomatitis virus (VSV; gbvrl) and human protein-coding genes were obtained from the Codon Usage Database (http://www.kazusa.or.jp/codon/), as described previously ^42^. Codon usage was expressed as frequency per 1,000 codons. Amino acids encoded by a single codon (e.g., AUG for methionine and UGG for tryptophan), as well as stop codons, were excluded from subsequent analyses.

Codon usage was converted from the codon level to the tRNA level by summing the frequencies of synonymous codons decoded by each tRNA isoacceptor, according to the Percudani decoding rules. This conversion yielded a measure of total translational demand for each tRNA species. To quantify relative translational pressure, converted viral codon usage was normalized to the corresponding converted human codon usage and expressed as a log₂ ratio for each tRNA species.

### Polysome profiling

Polysome profiling was performed as previously described ^76^. Briefly, cells were treated as described in ‘Cell Culture and Chemical Treatments’ subsection, and incubated with cycloheximide (CHX) to a final concentration of 100 ug/mL for 3 minutes at 37°C. Cells washed twice with ice-cold PBS supplemented with 100 ug/mL CHX prior to lysis.

#### MCF7

*Cells were lysed* in 300 µL hypotonic lysis buffer (LB Immagina; cat. no. IBT0031) supplemented to final concentrations of 1% sodium deoxycholate, 5 U/mL DNase I, and 200 U/mL RiboLock RNase Inhibitor (Thermo, #EO0382). Lysates were clarified at 20,000xg for 5 min at 4°C to remove cell debris and nuclei. A total of 250 µL of cleared lysate was loaded onto a linear 15–50% sucrose (m/v) gradient prepared in 100 mM NaCl, 10 mM MgCl₂, and 10 mM Tris-HCl (pH 7.5). Gradients were subjected to ultracentrifugation using a Beckman SW41Ti rotor at 40,000 rpm for 1 h 40 min at 4 °C in a Beckman Optima LE-80K Ultracentrifuge.

#### HEK293T

Cells were lysed in 500 µL lysis buffer containing 100 mM KCl, 10 mM MgCl2, 20 mM Tris-HCl pH 7.4, 0.5% Triton X-100, 0.2M Sucrose, 2U/uL RNasein Ribonucleotide Inhibitor (Promega, #N211B), 0.1 mg/mL cycloheximide, 1 mM DTT and 1X EDTA-free protease inhibitor (Roche #11873580001). Lysates were clarified by centrifugation, and protein concentrations were determined using the Pierce™ BCA Protein Assay Kit (Thermo Fisher Scientific; cat. no. 23225) according to the manufacturer’s instructions. For each sample, 2 mg of cleared lysate was adjusted to an equal volume with lysis buffer and loaded onto a linear 10–50% (w/v) sucrose gradient prepared in gradient buffer containing 100 mM KCl, 10 mM MgCl₂, 20 mM Tris-HCl (pH 7.4), 0.1 mg mL⁻¹ cycloheximide, and 0.5 mM DTT. Gradients were centrifuged in a Beckman SW41Ti rotor at 35,000 rpm for 2 h 30 min at 4 °C using a Beckman Optima XPN-100 ultracentrifuge. Following ultracentrifugation, gradients were analysed by continuous monitoring of absorbance at 254 nm using an ISCO UA-6 UV detector (MCF7) or a Biocomp Gradient Station (HEK293T).

### Analysis of tRNAs from polysome fractions

Sucrose gradients were fractionated into twenty 1 mL fractions while continuously monitoring absorbance at 254 nm using a Biocomp Gradient Station. Fractions corresponding to heavy polysomes (fractions 13–20) were pooled and diluted to a final volume of 7 mL in polysome extraction buffer (PEB; 20 mM Tris-HCl pH 7.4, 100 mM KCl, 10 mM MgCl₂). The diluted sample was loaded onto a pre-rinsed Amicon Ultra-15 centrifugal filter unit (100 kDa cutoff; Millipore, cat. no. UFC910024) and centrifuged at 2,160xg for 20 min at 4°C. The retentate (<200 µL) was collected for RNA extraction. For RNA isolation, 500 µL TRIzol LS reagent (Thermo Fisher Scientific; cat. no. 10296010) was added to the retentate, vortexed, and incubated for 5 min at room temperature, followed by the addition of 100 µL chloroform. Samples were vortexed, incubated for an additional 5 min at room temperature, and centrifuged at 12,000xg for 5 min at 4°C. The aqueous phase was transferred to a fresh RNase-free tube, mixed with an equal volume of isopropanol and 1 µL Pellet Paint co-precipitant (Novagen; cat. no. 69049-3), and incubated overnight at −80°C. RNA was pelleted by centrifugation at 12,000xg for 10 min at 4°C, washed twice with 75% ethanol, and resuspended in RNase-free water. Samples were then incubated at 65°C for 5 min to facilitate resuspension. RNA integrity was assessed using TapeStation analysis, and RNA concentration was measured by NanoDrop. Size-based small RNA enrichment was not performed due to the low abundance of short RNA species in polysome-enriched fractions. Total tRNA isolated from heavy polysomes was directly subjected to nano-tRNA-seq library preparation.

### Quantification of polysome profiles

Polysome profiles were exported as position–absorbance tables. For each profile, the 80S monosome and polysome regions were defined in positional units along the gradient. The area under the curve (AUC) for the 80S peak and the polysomal region was calculated in R using trapezoidal numerical integration (‘*trapz’* function, pracma library version 2.4.4). For each biological replicate and condition, a translational efficiency index (TE) was defined as the ratio of polysomal AUC to 80S AUC (TE = AUC_poly / AUC_80S). These TE values from independent gradients were used for statistical analysis (two-sided unpaired Welch’s t-test) and are reported in **Fig. S5B,C**.

### Puromycin incorporation assays

Puromycin incorporation was performed using the Surface Sensing of Translation (SunSET) method^77^ to assess global protein synthesis. HEK293T cells were seeded in 6-well plates and cultured to approximately 70% confluency. Cells were washed with pre-warmed PBS and treated with puromycin (Merck, cat. no: P8833-25MG) at a final concentration of 0.5 μg/mL for 10 minutes at 37 °C. Immediately following treatment, cells were washed twice with ice-cold PBS and lysed in RIPA buffer [50 mM Tris-HCl (pH 7.5), 150 mM NaCl, 1% Triton X-100, 0.1% SDS, 0.5% sodium deoxycholate] supplemented with protease inhibitor (Roche cOmplete™ Protease Inhibitor Cocktail used at 1× final concentration, cat. No: #11873580001). Lysates were additionally treated with Turbo DNase (Thermo Fisher, cat. no: AM2239) at 2 U per sample for 1 hour on ice. Following lysis, samples were centrifuged at maximum speed (≥14,000 × g) for 5 minutes at 4 °C to remove debris. Protein concentrations were determined using the Pierce™ BCA Protein Assay Kit (Thermo Fisher, cat. no: 23225) according to the manufacturer’s instructions. Equal amounts of protein were resolved by SDS–PAGE and analyzed by Western blot using an anti-puromycin primary antibody (DSHB, cat. no: PMY-2A4) and an HRP-conjugated anti-mouse secondary antibody (Sigma-Aldrich, cat. no: #31430). Lane intensity was quantified using the iBright Analysis Software version 5.2.2 (Thermo Fisher Scientific) using identically sized rectangular regions of interest (ROIs) drawn over each lane. The puromycin signal was extracted as the local background–corrected volume for each ROI. Total protein loading was quantified from the corresponding Ponceau S–stained membrane, using the same ROIs. For each lane, the puromycin signal was normalized to the matched Ponceau S signal. For each amino-acid treatment (arginine or leucine), normalized puromycin values were scaled to the mean of the corresponding control condition, which was set to 1.0 for presentation in **Fig. S5A**.

### Data analysis and statistics

*Principal Component Analysis (PCA)*: PCA was performed on tRNA-Seq data using a custom R script (see ‘Data availability’ section). PCA on Ribo-Seq and RNA-seq was conducted on per-transcript read counts, modeled with a one-factor model ‘*y ∼ Treatment’* (Control and Treatment groups n=3), keeping top 10% differentially expressed genes between conditions. *Amino-acid percentage calculation*: Codon counts were aggregated based on their corresponding amino-acids. Amino-acid percentages were calculated for each sample as the ratio of amino acid count to total codon counts). *Correlation analysis*: Associations between variables were assessed using either Spearman’s rank correlation coefficient (rho) or Pearson’s correlation coefficient (r). These were used as estimates of effect size, quantifying the strength and direction of linear (Pearson) and non-linear (Spearman) relationships, respectively. Correlation coefficients and corresponding P-values were calculated using the ‘cor.test()’ function in R (method = ’pearson’ or ’spearman’).

### Per-site fragmentation analysis

After counting reads mapping to full-length tRNAs and tRNA fragments for each reference tRNA, one-factor differential expression analysis was performed (y ∼ Type, where Type = {Full-length, Fragment}) separately within each sample group (control, treatment, ribo-tRNA and total-tRNA) using the edgeR package^78^. Resulting p-values were corrected using the Benjamini-Hochberg method, and volcano plots were generated using the p-adjusted values and corresponding log2 fold changes (Fragment vs Full-length).

To identify the precise cleavage sites within tRNAs, a custom script ‘bams2ends.py’ was implemented (publicly available on GitHub: see *Data Availability*). Briefly, the analysis focused on the 5’ termini of aligned reads to detect cleavage hotspots. The 5′ termini of aligned reads from BAM files was extracted and aggregated at start positions by reference tRNA. Positions with sufficient read coverage were tested for significant enrichment of 5′ termini between conditions (Control vs. Treatment, Ribo-tRNA vs. total-tRNA) using edgeR’s negative binomial framework. P-values were corrected with the Benjamini–Hochberg method, and sites with significant differences in cleavage frequency were annotated as putative fragmentation loci relative to the tRNA reference coordinates.

### LC-MS/MS analysis of RNA modifications

100 ng of small RNA (<200 nt) fraction, separated as described above, were digested with 1 μl of the Nucleoside Digestion Mix (New England BioLabs, #M0649S) and the mixture was incubated at 37 °C for 1 h. Samples were analyzed using an Orbitrap Eclipse Tribrid mass spectrometer (Thermo Fisher Scientific, San Jose, CA, USA) coupled to an EASY-nLC 1200 (Thermo Fisher Scientific (Proxeon), Odense, Denmark). Ribonucleosides were loaded directly onto the analytical column and were separated by reversed-phase chromatography using a 50-cm column with an inner diameter of 75 μm, packed with 2 μm C18 particles (Thermo Fisher Scientific, cat # ES903). Chromatographic gradients started at 93% buffer A and 3% buffer B with a flow rate of 250 nl/min for 5 minutes and gradually increased to 30% buffer B and 70% buffer A in 20 min. After each analysis, the column was washed for 10min with 0% buffer A and 100% buffer B. Buffer A: 0.1% formic acid in water. Buffer B: 0.1% formic acid in 80% acetonitrile. The mass spectrometer was operated in positive ionization mode with nanospray voltage set at 2.4 kV and source temperature at 275°C. For the Parallel Reaction Monitoring (PRM) method, the quadrupole isolation window was set to 1.4 m/z, and MS/MS scans were collected over a mass range of m/z 50-450, with detection in the Orbitrap at the resolution of 60,000. MSMS fragmentation of defined masses with schedule retention time was performed using HCD at NCE 20, the auto gain control (AGC) was set to “Standard” and a maximum injection time of 118 ms was used. In each PRM cycle, one full MS scan at a resolution of 120,000 was acquired over a mass range of m/z 220-700 with detection in the Orbitrap mass analyzer. Auto gain control (AGC) was set to 1e5 and the maximum injection time was set to 50 ms. Serial dilutions were prepared using commercial pure ribonucleosides (0.005-150 pg, Carbosynth, Toronto Research Chemicals) in order to establish the linear range of quantification and the limit of detection of each compound. A mix of commercial ribonucleosides was injected before and after each batch of samples to assess instrument stability and to be used as an external standard to calibrate the retention time of each ribonucleoside. Acquired data were analyzed with the

Skyline-daily software (v24.1.1.254). Extracted precursor areas of the ribonucleosides were used for quantification, and absolute picogram amounts were derived from external calibration curves. To account for technical variability in ionization and injection volume, picogram values were normalized to the median AUGC (A, U, G, C base nucleosides) value for each sample. Normalization ensured comparability across conditions while preserving relative differences between nucleosides. The metabolomics data have been deposited to MetaboLights ^79^ repository with the study identifier MTBLS12806.

## DATA AVAILABILITY

Base-called FAST5 have been deposited in the European National Archive (ENA) under accession codes PRJEB89426, PRJEB106334, and PRJEB106349. Raw LC-MS/MS data have been deposited to the Metabolight repository with the dataset identifier MTBLS12806. MS data deposit accession found in **Table S17**. All sequencing runs used in this work are listed in **Table S18.** Processed data (tRNA counts) for all tRIBO-seq and Nano-tRNAseq experiments performed in this work are available in **Table S19**.

## CODE AVAILABILITY

All code used in this work is available at: https://github.com/novoalab/tRIBO-seq

## ACKNOWLEDGEMENTS

We thank Cristina Petrella, Xavier Hernandez Alias and Hannah Benisty for their guidance and support in establishing the starvation experiments. We also thank Leszek Pryszcz for his bioinformatic help with the per-site fragmentation analysis, and Alexane Ollivier for her assistance in preparing the HEK293T untreated libraries for sequencing. HY is supported by the European Union’s Horizon 2020 research and innovation programme under the Marie Skłodowska-Curie grant agreement (H2020-MSCA-ITN-2020 No 956810). MM is supported by the Juan de la Cierva fellowship (JDC2023-050553-I funded by MICIU/AEI /10.13039/501100011033 and FSE+). DR and AS are funded by FEDER (Fundo Europeu de Desenvolvimento Regional) through COMPETE 2030 (Project COMPETE2030-FEDER-00758600) and by Portuguese national funds through FCT (Fundação para a Ciência e a Tecnologia), under the iBiMED project UID/04501/2025. JS is funded by FCT PhD grant 2022.14102.BD. This work was supported by the European Union’s Horizon Europe through the European Research Council (ERC-StG-2021 No 101042103 to EMN). We acknowledge the support of the Spanish Ministry of Science and Innovation through the Centro de Excelencia Severo Ochoa (CEX2020-001049-S, MCIN/AEI /10.13039/501100011033), and the Generalitat de Catalunya through the CERCA programme. The proteomics analyses were performed in the CRG/UPF Proteomics Unit which is part of the Spanish National Infrastructure for Omics Technologies (ICTS OmicsTech).

## AUTHOR CONTRIBUTIONS

MM performed cell culture, treatment and preparation for all HEK293T experiments, interpreted data analyses from all experiments and contributed to experimental design. HY performed all bioinformatic analyses and curated the GitHub repository. ADP, MA and IB performed cell cultures, treatment and sample processing for the Arsenite experiment. JS, DR and ARS performed the cell cultures, treatment and sample processing for the VSV-infection experiment. ADP and MA performed cell cultures, treatment and preparation for Harringtonine and Methionine starvation experiments. LLL prepared the samples for LC-MS/MS and helped with tRIBO-seq libraries. EMN and MC conceived and supervised the project. MM and HY built the figures. HY, MM, MC and EMN wrote the paper, with contributions from all authors.

## DECLARATION OF INTERESTS

EMN has received travel and accommodation expenses to speak at Oxford Nanopore Technologies conferences. MM has received travel bursaries from ONT to present her results at research conferences. EMN is a member of the Scientific Advisory Board of IMMAGINA Biotechnology s.r.l. MC is chief executive officer and shareholder of IMMAGINA Biotechnology s.r.l.. EMN is listed as inventor in the patent describing the Nano-tRNAseq method (PCT/IB2023/059599, publication WO2024/069467), which is licensed to IMMAGINA Biotechnology. HY, MA, IB and ADP are employees of IMMAGINA Biotechnology s.r.l.

