## Supplementary Figure for "Selective profiling of translationally active tRNAs and their dynamics under stress"

** Equal contribution*

**Figure S1: Sequencing active tRNAs by co-eluting tRNAs from elongating ribosomes**. (**A**) Schematic overview of the tRIBO-seq workflow, illustrating the key steps in capturing and sequencing ribosome-associated tRNAs. A total of 9 independent biological samples from HEK293T cells are shown. (**B**) TapeStation RNA profiles showing total RNA from cell lysate samples and corresponding <200 nt fractions obtained after clean-up (see Methods). These samples were used as input for total-tRNA (Nano-tRNAseq) library preparation. **(C)** TapeStation RNA profiles of ribosome-captured RNAs, used as input for ribo-tRNAs (tRIBO-seq) library preparation. Ribosome-bound RNA samples exhibit a marked enrichment of ribosomal RNA (rRNA) and tRNAs compared to total input RNA, highlighting selective recovery of actively engaged RNAs. Rec.: Recommended. As the adaptors selectively capture tRNAs, the clean-up step is optional when material is limited. (**D**) Barplots showing the mean RNA recovery (in µg) at each step of the workflow, for both total tRNA and ribosome-associated tRNA (ribo-tRNA) samples. Error bars indicate standard error of the mean. (**E)** Stacked barplots depicting the relative proportion of mapped reads to cytoplasmic tRNA (cyt-tRNA), mitochondrial tRNA (mito-tRNA), and rRNA.

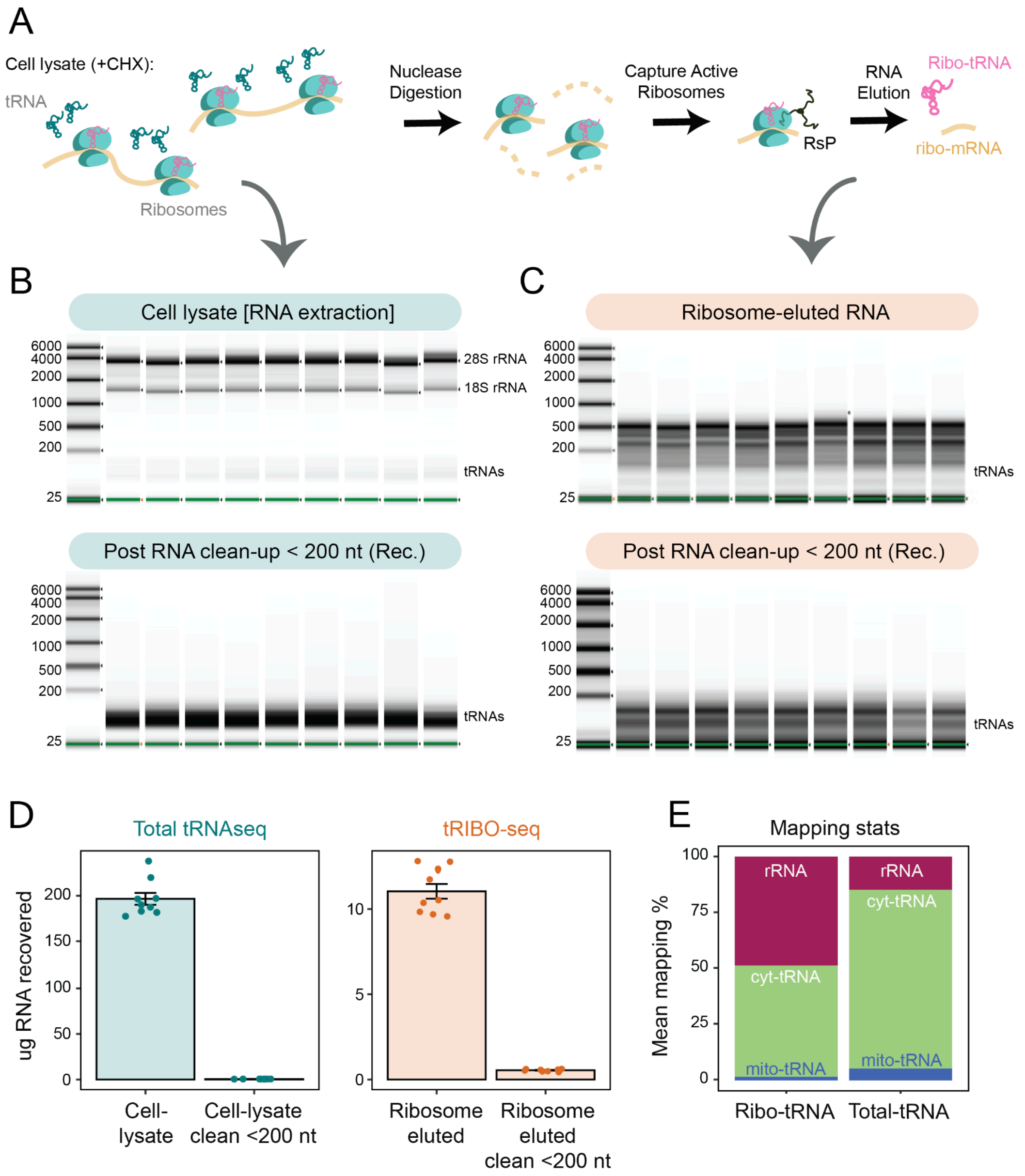

**Figure S2: Benchmarking tRIBO-seq against polysome-derived tRNA sequencing (Poly-seq). (A)** Schematic comparison of experimental workflows for tRIBO-seq (left) and polysome profiling followed by tRNA sequencing (right). While tRIBO-seq relies on a rapid ribosome pulldown, polysome profiling requires specialised equipment (shown in italics) and involves longer processing times. RNA recovery yields are indicated as the ratio of recovered RNA to input RNA, see **Table S1**. **(B)** Representative polysome profile from HEK293T cells showing absorbance at 260 nm across the sucrose gradient; the dashed line indicates fractions corresponding to heavy polysomes used for downstream tRNA sequencing. **(C)** TapeStation profiles of RNA recovered from heavy polysome fractions. **(D)** Reproducibility of polysome-derived tRNA sequencing (Poly-seq) and tRIBO-seq across biological replicates, assessed by pairwise correlation of tRNA abundance profiles (Pearson's R shown). **(E)** Principal component analysis (left) of tRNA abundances obtained by tRIBO-seq and polysome sequencing. **(F)** Correlation of mean log2(counts) in tRNA abundance between the two methods (right). Initiator methionine (iMet-CAT) and selenocysteine (SeC-TCA) tRNA is highlighted (n = 3 Poly-seq replicates, n = 4 tRIBO-seq replicates). Underlying data is reported in **Table S3**.

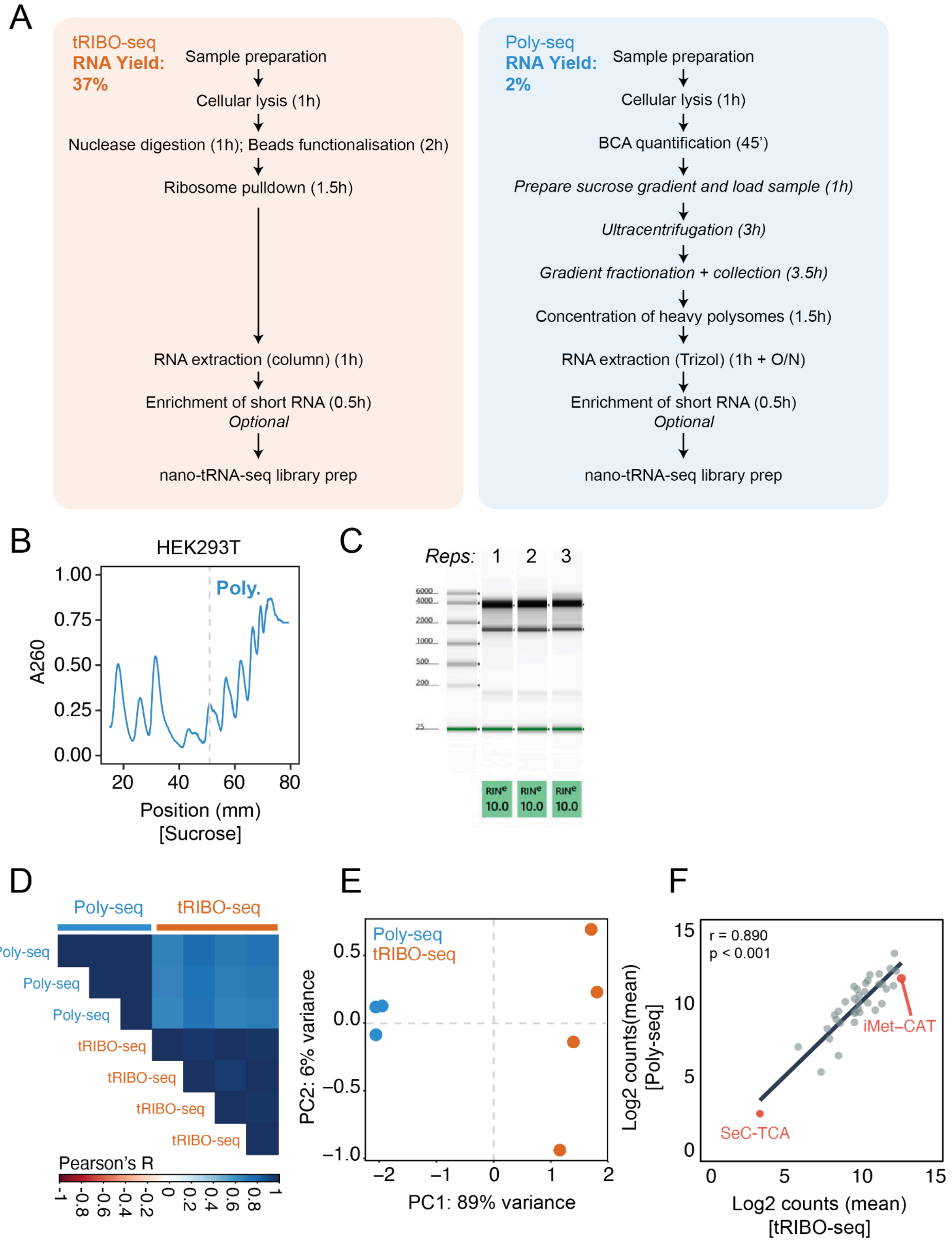

**Figure S3: Differential modification between total- and ribo-tRNAs. (A)** Heatmap of the differential basecalling errors of Ribo-tRNAs relative to Total-tRNAs. The x-axis represents nucleotide positions along the tRNA (5' to 3'). The y-axis corresponds to different tRNA isoacceptors (alphabetically ordered). Differential base-calling errors can be used as a proxy for differential modifications, as previously described [1]. (**B**) Bar plots for ribo-tRNAs (top) and total-tRNAs (bottom) showing the percentage of deletions at the CCA-tail positions (C1,C2,A). These provide a position-resolved visualization of differences at the 3′ end. Data are from HEK293T cell cultures, with *n = 2* independent biological replicates per condition. We should note that the high adaptor-to-tRNA ratio used both in tRIBO-seq and Nano-tRNAseq library preparation enables the capture of incomplete tRNAs, including those lacking full CCA tails, as previously shown [1].

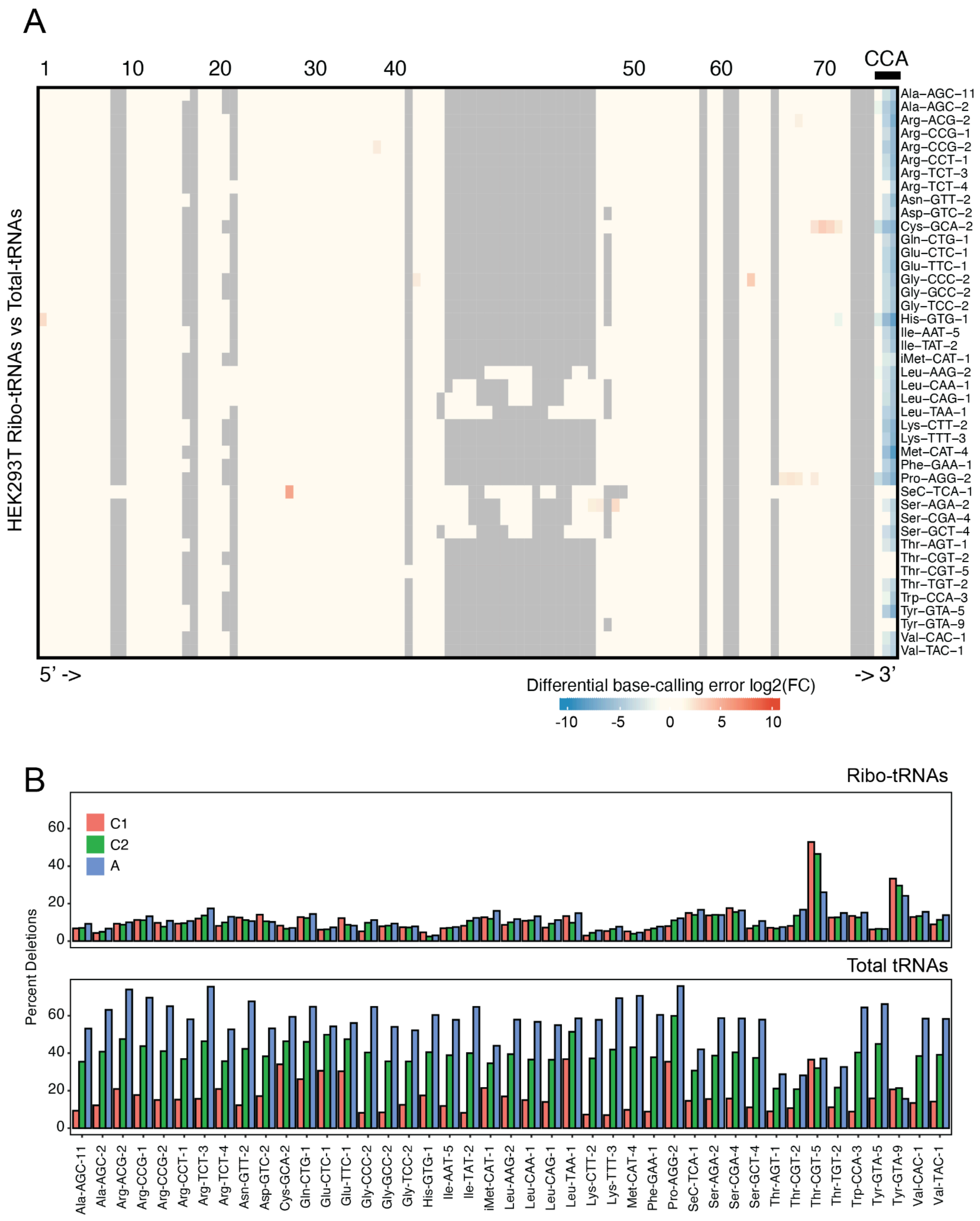

**Figure S4: Harringtonine experiment confirms the specificity of tRIBO-seq at capturing ribo-embedded tRNAs. (A)** Schematic representation of HAR mechanism of translational block. Briefly, translation initiation begins when the 43S pre-initiation complex (PIC)- comprising the 40S ribosomal subunit, initiation factors, and the eIF2–GTP–Met-tRNA$_i$ ternary complex- associates with the eIF4 complex bound to the 5′ cap of the mRNA. The PIC then scans the mRNA until encountering the start codon, where 60S subunit joining produces the 80S initiation complex and elongation commences. Harringtonine (HAR) inhibits elongation by blocking the A-site of newly formed 80S ribosomes, causing ribosomes to accumulate at start codons while preventing further elongation. For a detailed description of HAR's mechanism, see[2]. **(B)** Fold-change (FC) difference of tRNA abundances between HAR vs CTRL conditions for tRIBO-seq (orange) and total-tRNA-seq (teal). iMet-CAT (shown in red) is significantly enriched in tRIBO-seq following HAR treatment, consistent with ribosome stalling at initiation sites. **(C)** Scatterplots depicting the correlation between ribo-tRNA and total-tRNA abundances, for control conditions. The Pearson $R^2$ correlation is also shown. Error bars indicate standard error of the mean. **(D)** Sample of replicate-wise scatterplots comparing log-normalized tRNA abundances for biological replicate pairs under control (left), HAR (middle), and HAR vs. CTRL conditions (right). The strong correlation across replicates further confirms the robustness and reproducibility of the tRIBO-seq measurements. **(E)** Correlation matrix showing pairwise Pearson correlation coefficients between the three biological replicates for both tRIBO-seq and total-tRNA-seq under HAR and control conditions. High replicate concordance indicates strong reproducibility of the tRNA abundance profiles. **(F)** Scatterplots showing the correlation between mean codon abundance at the ribosomal P-site in control cells collapsed by amino-acid (y-axis) with tRNA abundances, using either ribo-tRNAs (left) or total tRNAs (right) populations (x-axis), respectively (n = 20). Spearman's rho and associated p-values are also shown. All experiments were performed using MCF-7 cell cultures, with *n* = 3 independent biological replicates per condition.

*(figure in next page)*

**Figure S4** (legend in previous page)

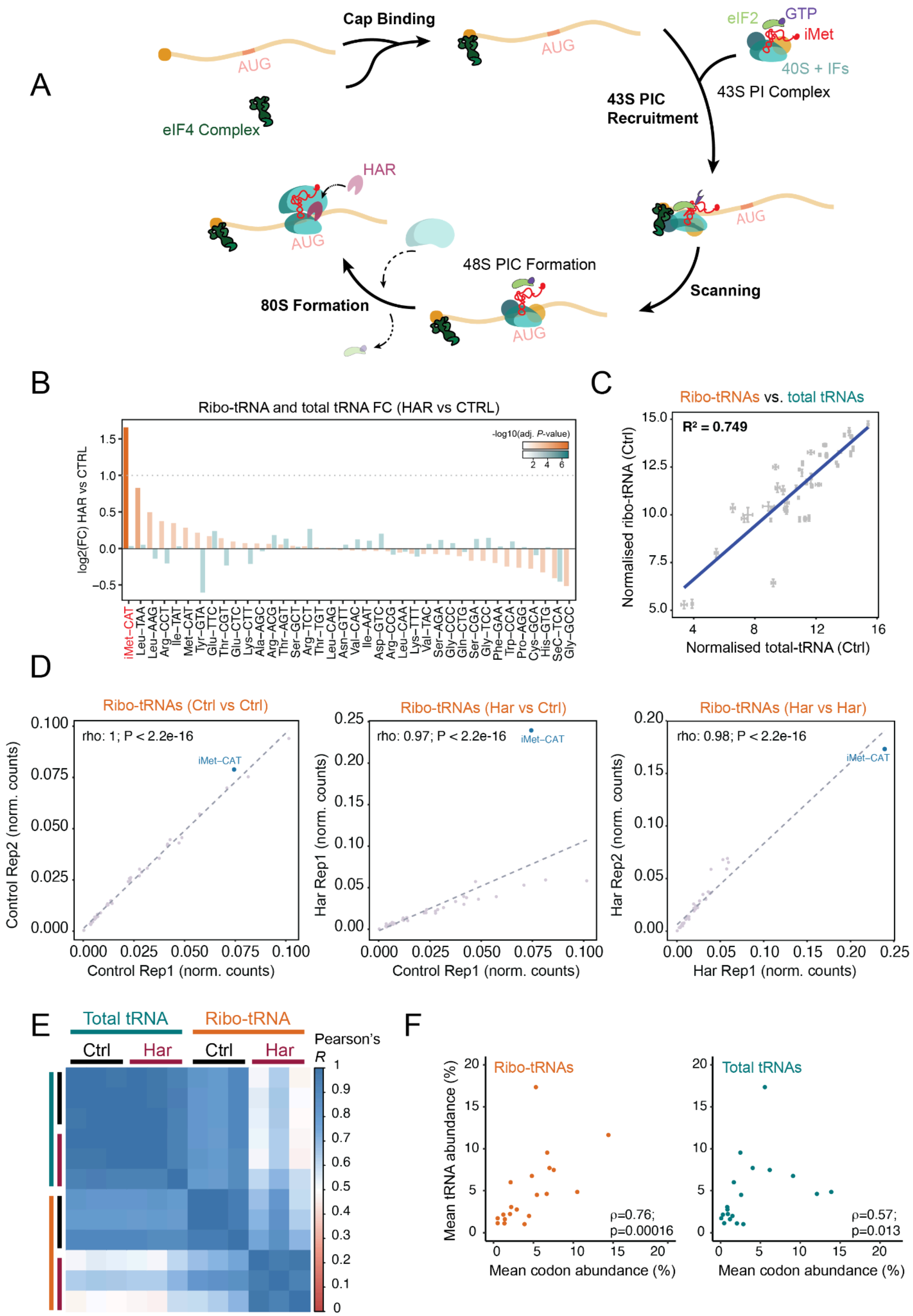

**Figure S5: Validation of amino acid deprivation effects on cellular translation activity. (A)** Puromycin incorporation assay. Upper panel: Ponceau S staining was used as protein loading control. Lower panel: Western blot of total protein lysates from HEK293T cells treated with amino acid starvation for 3 h or 6 h compared to control (Ctrl), probed with anti-puromycin antibody. Right: Quantification of puromycin signal normalized to corresponding Ponceau S signal (see *Methods*), showing a reduction in global protein synthesis after 3 h and 6 h of starvation. Error bars represent standard deviation (n = 2). Two-sided unpaired Welch's t-test results are reported. **(B,C)** Left panels: Representative polysome traces for Arg-deprived versus control conditions (above) and leucine-deprived versus control conditions (lower). Monosome (M, red-shaded) and polysome (Poly., green-shaded) regions indicated. Right panels: Quantification of ratio of polysome-associated area under the curve (AUC) to 80S monosome AUC shows a significant reduction in active translation under starvation conditions (3h) compared to controls. Error bars represent standard deviation (n = 2); two-sided unpaired Welch's t-test results reported. Statistical significance is annotated as follows: n.s.: non significant, *$P < 0.05$, **$P < 0.01$, ***$P < 0.005$.

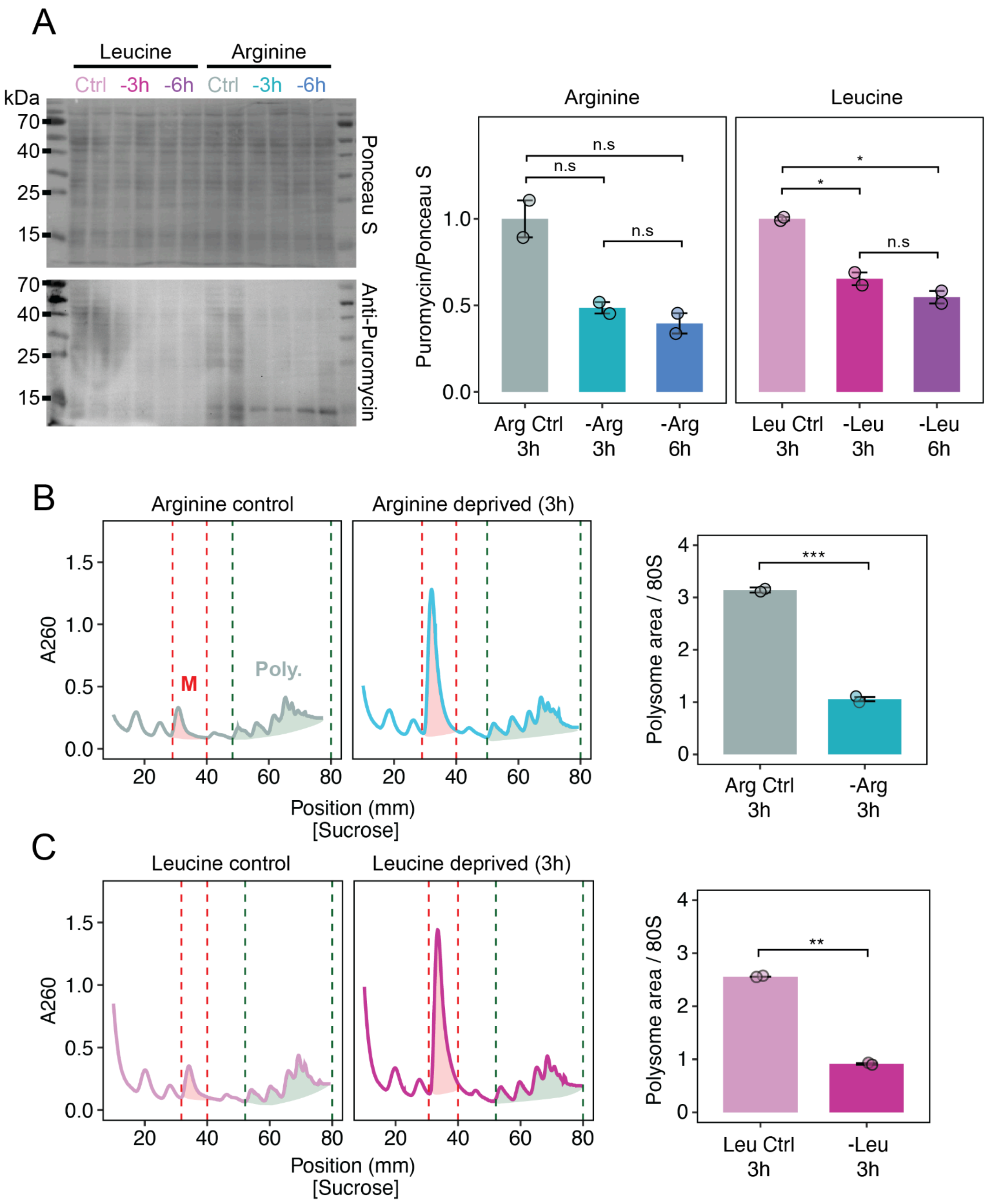

**Figure S6: Ribosome profiling of HEK cells upon 3-hour Leucine deprivation. (A)** PCA of Ribo-seq data from Leucine-deprived and controls cells at 3 hours. Each condition was performed with $n$ = 3 independent biological replicates. **(B)** Distribution of ribosome-protected fragment (RPF) lengths after processing; bars show mean +/- s.d. across biological replicates ($n$ = 3). **(C)** Proportion of processed RPFs assigned to genomic regions and reading frames. **(D)** Metagene profiles showing average ribosome occupancy across transcripts, aligned to the start and stop codons. (E) Volcano plots summarizing differential enrichment of codons inferred at the E-, P-, and A-sites. Each point corresponds to one codon; the x-axis shows log2 fold change (leucine deprivation vs control). The dashed horizontal line indicates the significance threshold (p-adj < 0.05). Leucine codons are highlighted and labeled in orange, and show a pronounced enrichment at the A-site, consistent with ribosome pausing when leucine is limiting.

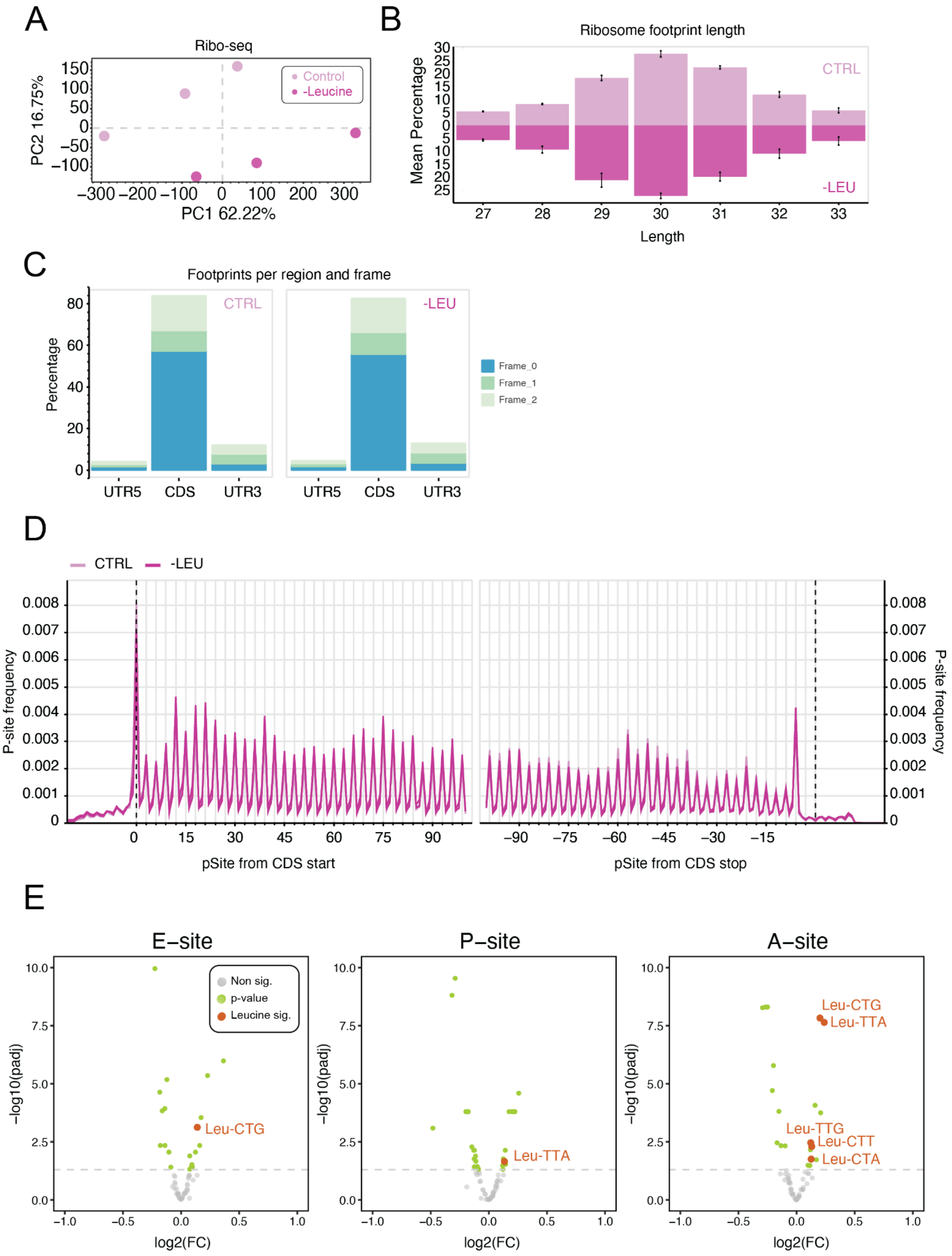

**Figure S7: Extended differential expression analysis between amino-acid deprivation conditions**. **(A)** PCA of total tRNAs between experimental conditions. Each condition was performed with *n* = 3 independent biological replicates. **(B)** Volcano plots showing differential abundance of tRNAs between ribo-tRNAs and total-tRNAs for each condition. Positive fold change indicates more in ribo-tRNAs. For the volcano plot, significance was defined as $p < 0.05$ with an absolute log2 fold-change ≥ 1. Full-length tRNA reads were used for pairwise analysis (see *Methods*).

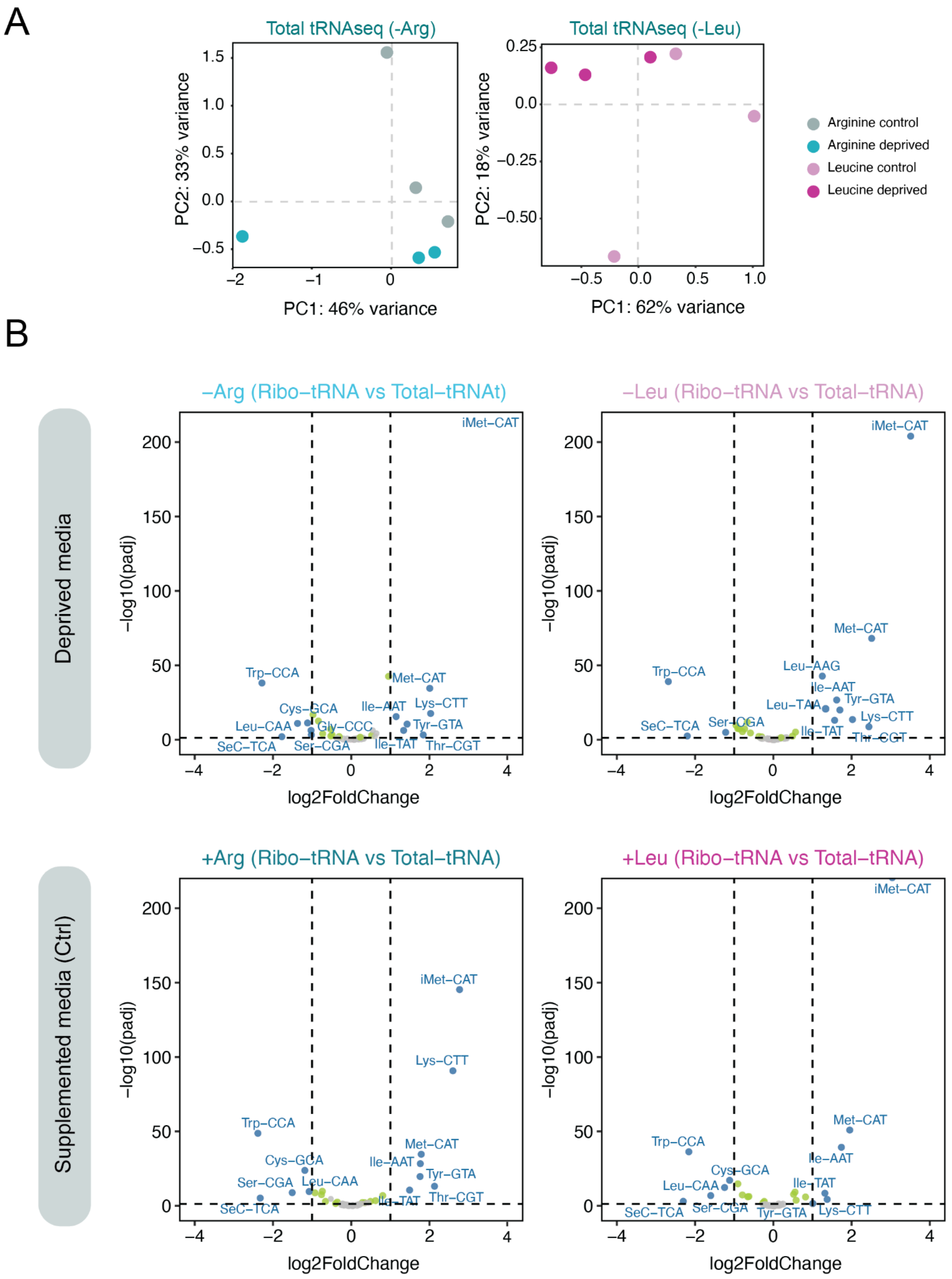

**Figure S8: Time-dependent response of ribosome-associated and total tRNAs to arginine deprivation. (A-B)** Volcano plots showing differential tRNA abundance in ribo-tRNAs (left) and total tRNAs (right) after 6h (top row) and 16h (bottom row) of arginine deprivation. For the volcano plot, significance was defined as $p < 0.05$ with an absolute log2 fold-change ≥ 1. No full-length filtering was applied in the pairwise analysis, as extended deprivation times resulted in low numbers of full-length tRNA reads (**Table S10**). **(C-D)** PCA plots for ribo-tRNAs (left) and total tRNA (right) shown for 6h (top row) and 16h (bottom row). Data represent n = 3 biological replicates per condition.

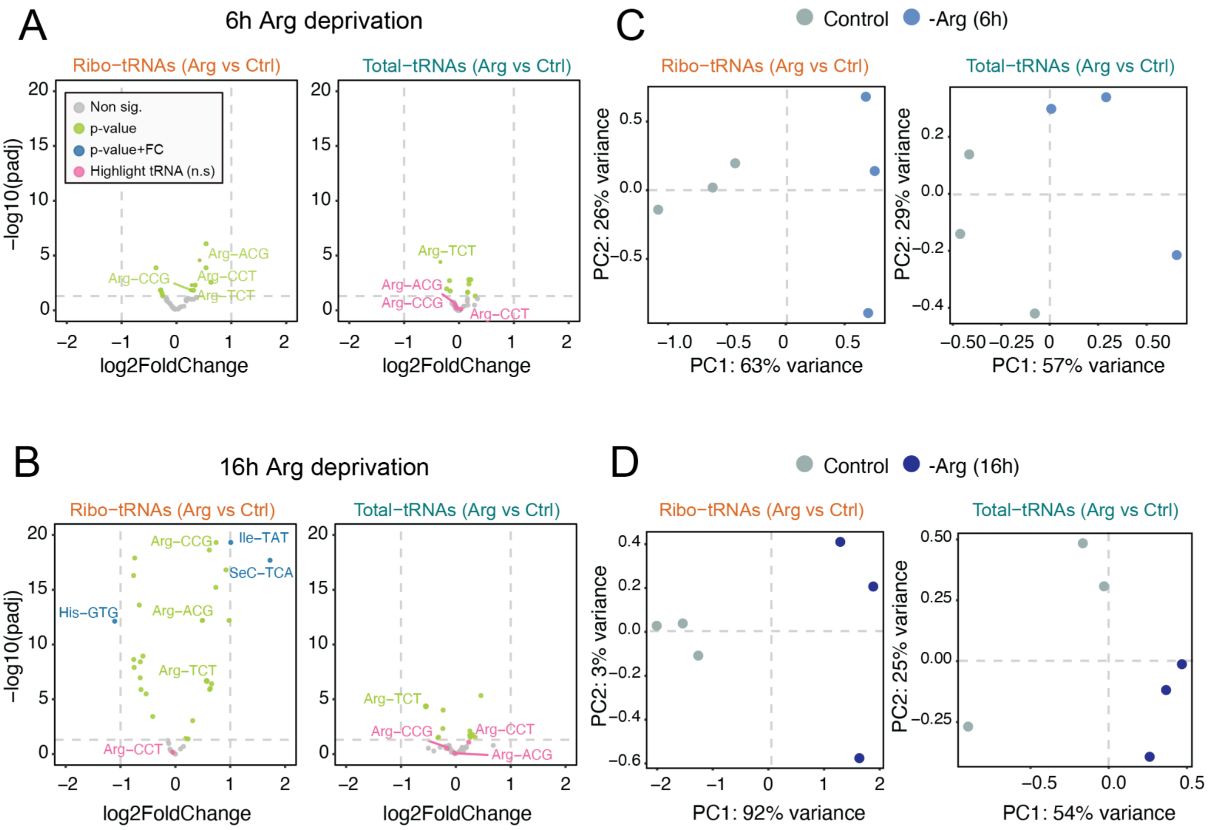

**Figure S9: Validation of transcriptional response upon methionine starvation. (A)** PCA of RNA-seq data from methionine-starved and control cells at 6 hours (left) and 16 hours (right), performed using the top 10% most significantly changed genes. Each condition was performed with n = 3 independent biological replicates. **(B)** Pairwise Pearson correlation heatmap of normalized gene expression across samples (n = 3 biological replicates per condition/timepoint). Values have been rounded to keep only two decimals. **(C)** Volcano plots showing differential RNA expression methionine starved at control conditions at 6h (left) and 16 hours (right). For the volcano plot, significance was defined as $p < 0.05$ with an absolute log2 fold-change ≥ 1.

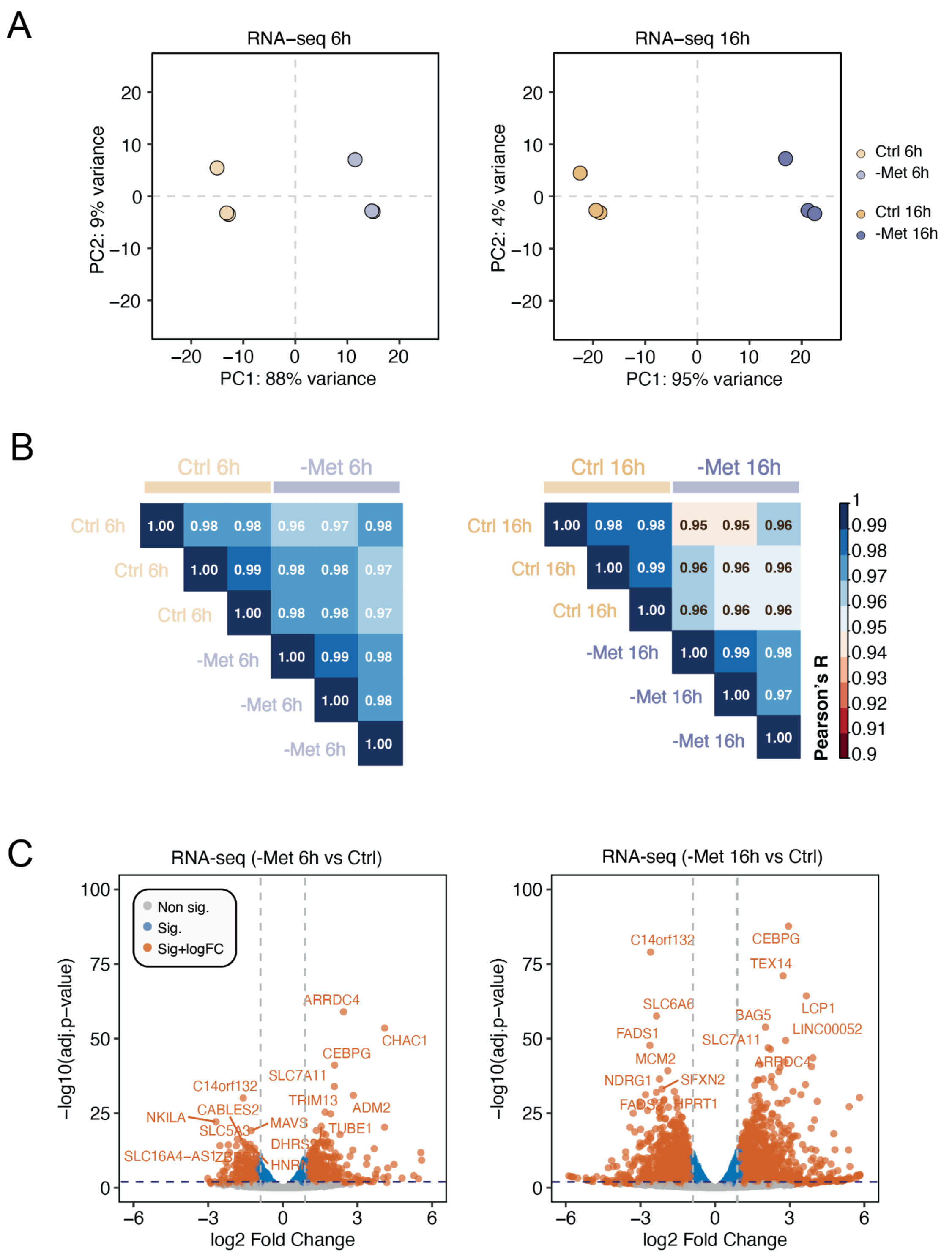

**Figure S10. Ribosome profiling quality control plots of methionine starvation experiments. (A)** PCA of Ribo-seq data from methionine-starved and control cells at 6 hours (left) and 16 hours (right), performed using the top 10% most significantly changed genes.. Each condition was performed with *n* = 3 independent biological replicates. **(B)** Ribosome footprint length distribution post-processing for methionine starved and control samples. Bars represent mean ± s.d. for *n* = 3 biological replicates. **(C)** Stacked barplot classifying processed footprints by frame position and genomic region. **(D)** Metagene plots showing ribosome occupancy for samples treated for 6h (upper panel) and 16h (lower panel). **(E)** Volcano plots summarizing differential enrichment of codons inferred at the A-sites for 6h (left) and 16h (right) methionine deprivation. Each point corresponds to one codon; the x-axis shows log2 fold change (deprivation vs control). The dashed horizontal line indicates the significance threshold (p-adj < 0.05). Methionine (Met-CAT) is highlighted and labeled in orange.. Ribo-seq data found at **Table S11**.

*(figure in next page)*

**Figure S10** (legend in previous page)

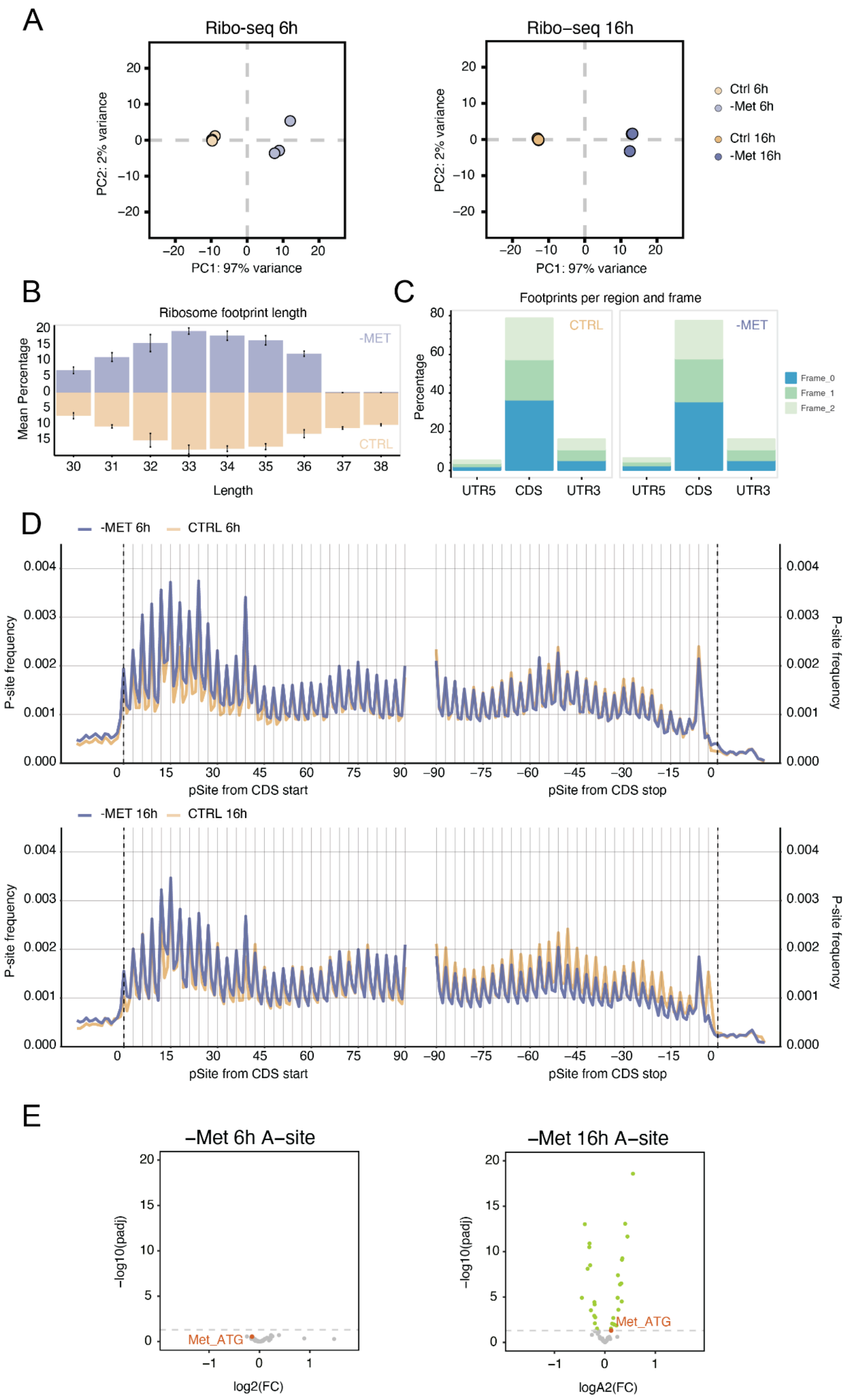

**Figure S11: S-adenosylmethionine (SAM) metabolism in MCF-7 cells. (A)** Biochemical map of key metabolic pathways related to SAM biosynthesis and utilization, highlighting links between methionine, arginine, glycine, serine, and threonine metabolism. MCF-7 cells lack MTAP, a key enzyme in the methionine salvage pathway; this is indicated by a crossed-out arrow. Schematic adapted from Tang *et al* [3]. **(B)** Intron retention in the *MAT2A* transcript observed under methionine starvation, leading to increased *MAT2A* expression. This response is driven by METTL16, a SAM-sensing methyltransferase that promotes *MAT2A* mRNA splicing and maturation under low SAM conditions [4]. **(C)** Expression changes in *MAT2A* gene under methionine starvation compared to control, assessed by RNA-seq and Ribo-seq at 6h and 16h. Bars represent mean normalized abundance ± s.d. ($n$ = 3 independent biological replicates per condition**)**. Differential expression was determined using the DESeq2 framework (see *Methods*). Adjusted p-values (Benjamini–Hochberg corrected) are indicated: RNA-seq 6 h FDR = 0.02; RNA-seq 16 h FDR = 0.04; Ribo-seq 6 h FDR = $4.1 \times 10^{-15}$; Ribo-seq 16 h FDR = $6.6 \times 10^{-40}$. Asterisks denote significance thresholds: **FDR < 0.05, **FDR < 0.01. ***FDR < 0.001.*

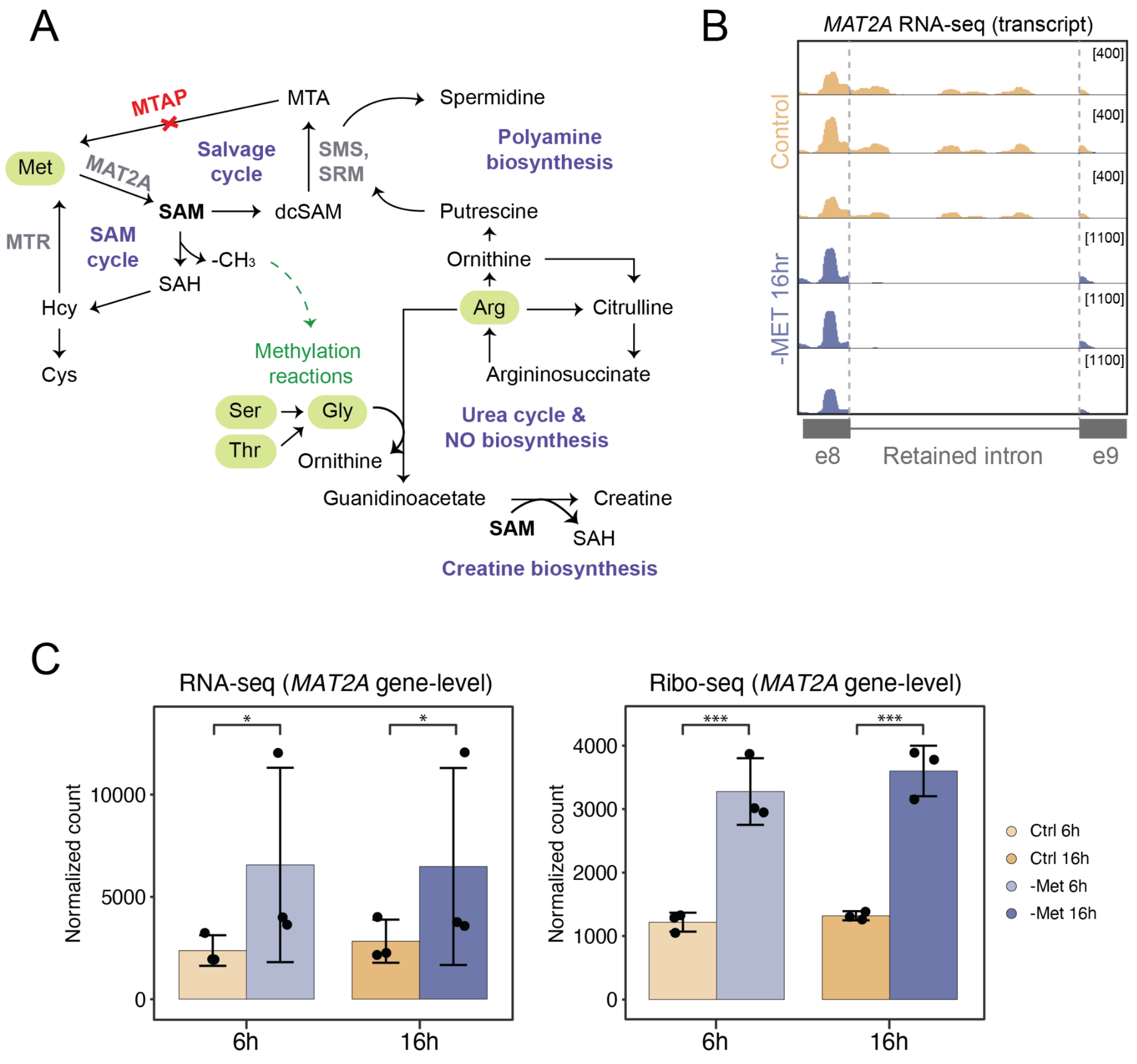

**Figure S12: Extended analysis of tRNA profiles upon methionine starvation. (A)** PCA of total tRNAs and Ribo-tRNAs across experimental conditions (*n* = 3 independent biological replicates per condition**)**. Ribo-tRNAs show clearer separation between methionine-starved and control samples at 16 hours, but not at 6 hours. Differential basecalling errors are used as a proxy to quantify differential modifications across two conditions [1]. **(B)** Heatmaps showing differential base-calling error rates (methionine-starved vs. control) for Ribo-tRNAs (top) and total tRNAs (bottom) at 6 hours. The x-axis indicates nucleotide position; the y-axis shows tRNA isoacceptors (alphabetically ordered).

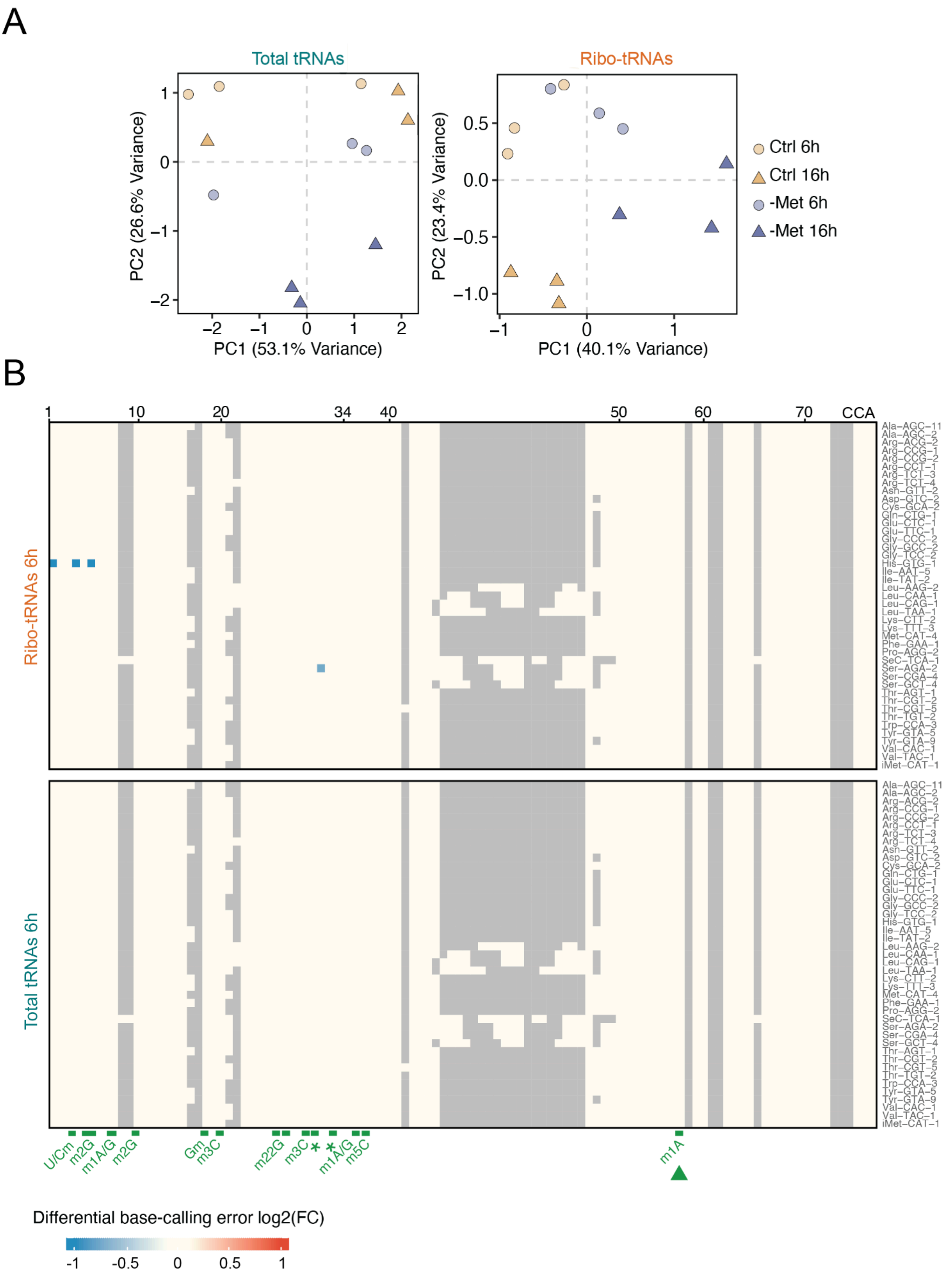

**Figure S13: Arsenite triggers the integrated stress response (ISR) pathway**. **(A)** Schematic representation of how arsenite induces reactive oxygen species (ROS) production via both mitochondria and NADPH oxidase (Nox), ultimately leading to ISR activation. **(B)** Left: Volcano plot showing differential gene expression based on ribosome footprints between arsenite and control conditions. Top five up- and downregulated genes are labeled. Right: the equivalent for matched RNA-seq data, here key ISR-related transcripts highlighted in bold (*n* = 3 independent biological replicates per condition**).** Significance was defined as $p < 0.05$ with an absolute log2 fold-change ≥ 1. See also **Table S12.**

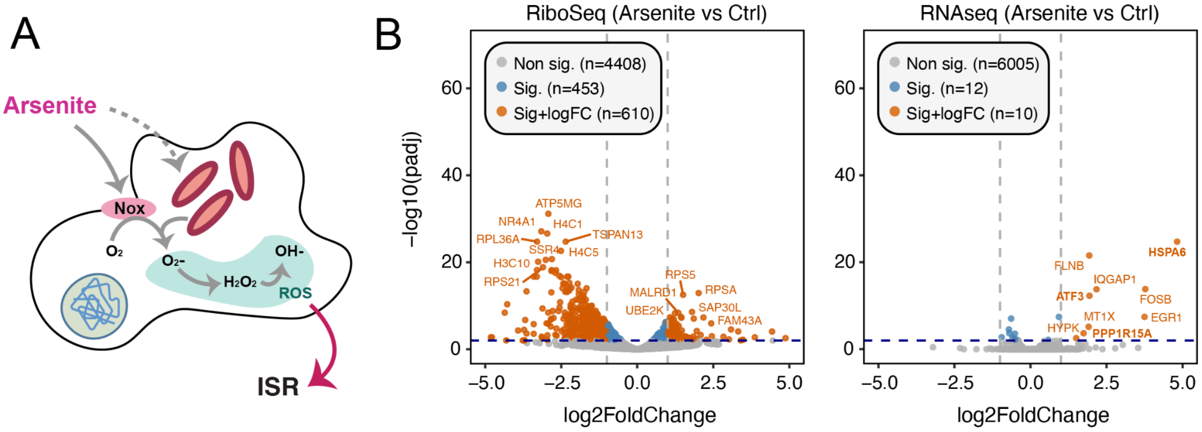

**Figure S14: Comparison of translational efficiency (TE) regulation under different stress conditions.** Scatterplots show $\log_2$ fold changes in ribosome occupancy (x-axis) versus RNA abundance (y-axis), derived from Ribo-seq and RNA-seq, respectively. Each gene is coloured based on its transcriptional and translational response, into 5 different categories: i) transcription: genes with significant transcriptome-only differences; ii) translation: genes with significant translation-only differences; iii) homodirection: genes with significant different both at transcription and translational level; iv) crossdirection: genes with differences in both transcription and translation but distinct up-/down-regulation directions; and v) unchanged: genes with no significant differences neither at transcriptional nor translational level. Barplots to the right indicate the number of genes per category. **(A)** Response to arsenite stress, with top regulated genes annotated. **(B)** Response to methionine deprivation. Each condition was performed with $n$ = 3 independent biological replicates. Ribo-seq data found at **Table S11** (methionine starved) and **Table S14** (arsenite treatment).

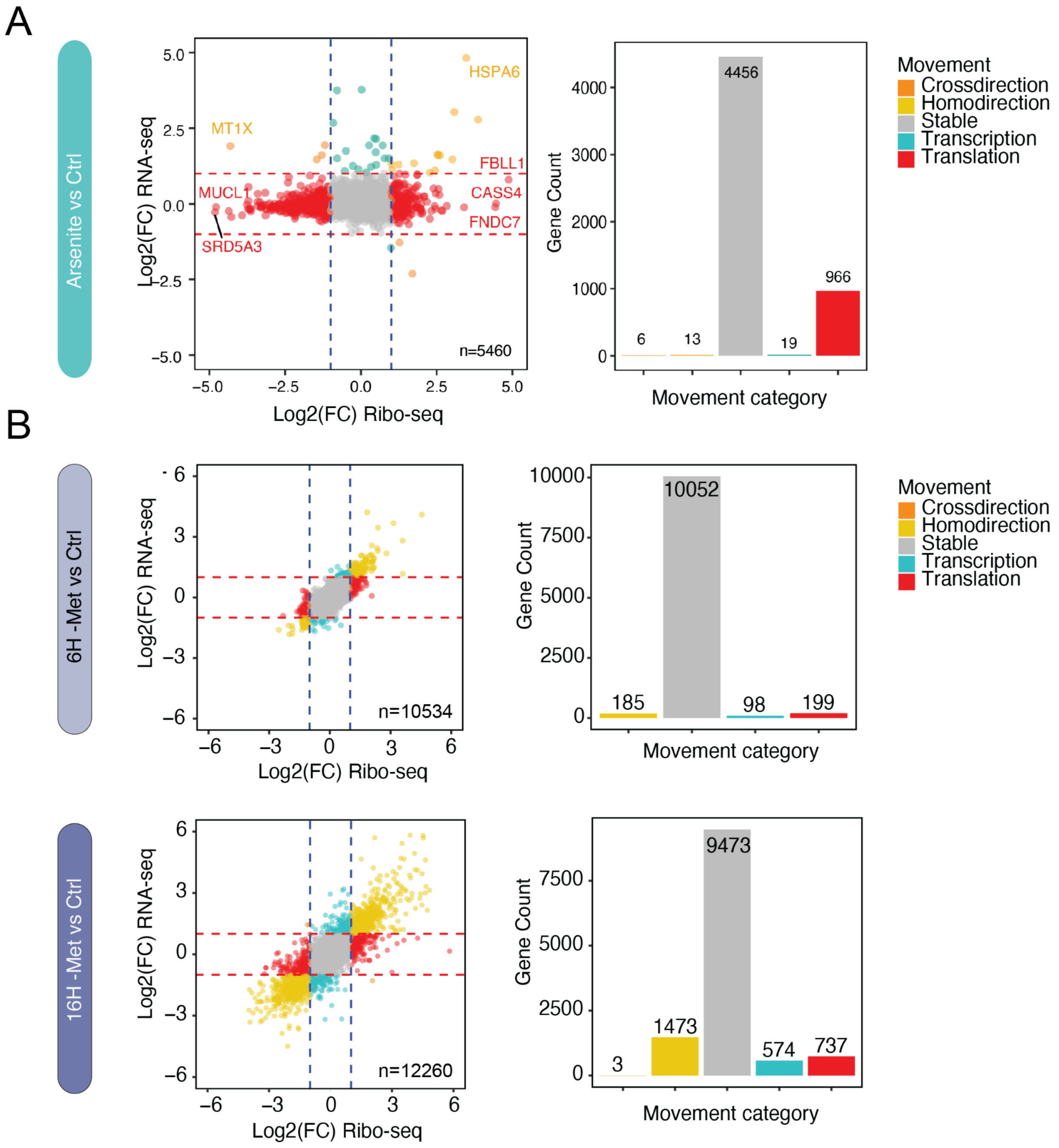

**Figure S15: Arsenite treatment affects tRNA dynamics at the amino acid (isoacceptor) level.** Volcano plots for each pairwise comparison of tRNA abundances, aggregated at the isoacceptor level (i.e., by amino acid identity rather than anticodon sequence, n = 20). Each condition was performed with $n$ = 3 independent biological replicates. Significance was defined as $p < 0.05$ with an absolute log2 fold-change ≥ 1.

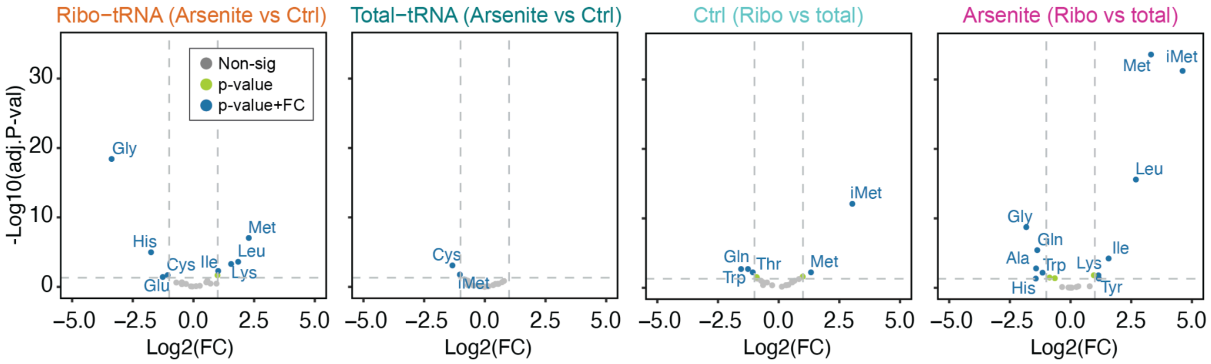

**Figure S16: Quality control plots for ribosome profiling of arsenite-treated samples. (A)** PCA of Ribo-seq data from arsenite-treated and control cells. **(B)** Distribution of ribosome footprint lengths after preprocessing in methionine-starved and control samples. Bars represent mean ± s.d. for *n* = 3 biological replicates. **(C)** Stacked bar plot classifying processed ribosome footprints by reading frame and genomic region**. (D)** Metagene plots showing average ribosome occupancy profiles for arsenite-treated and control samples (*n* = 3).

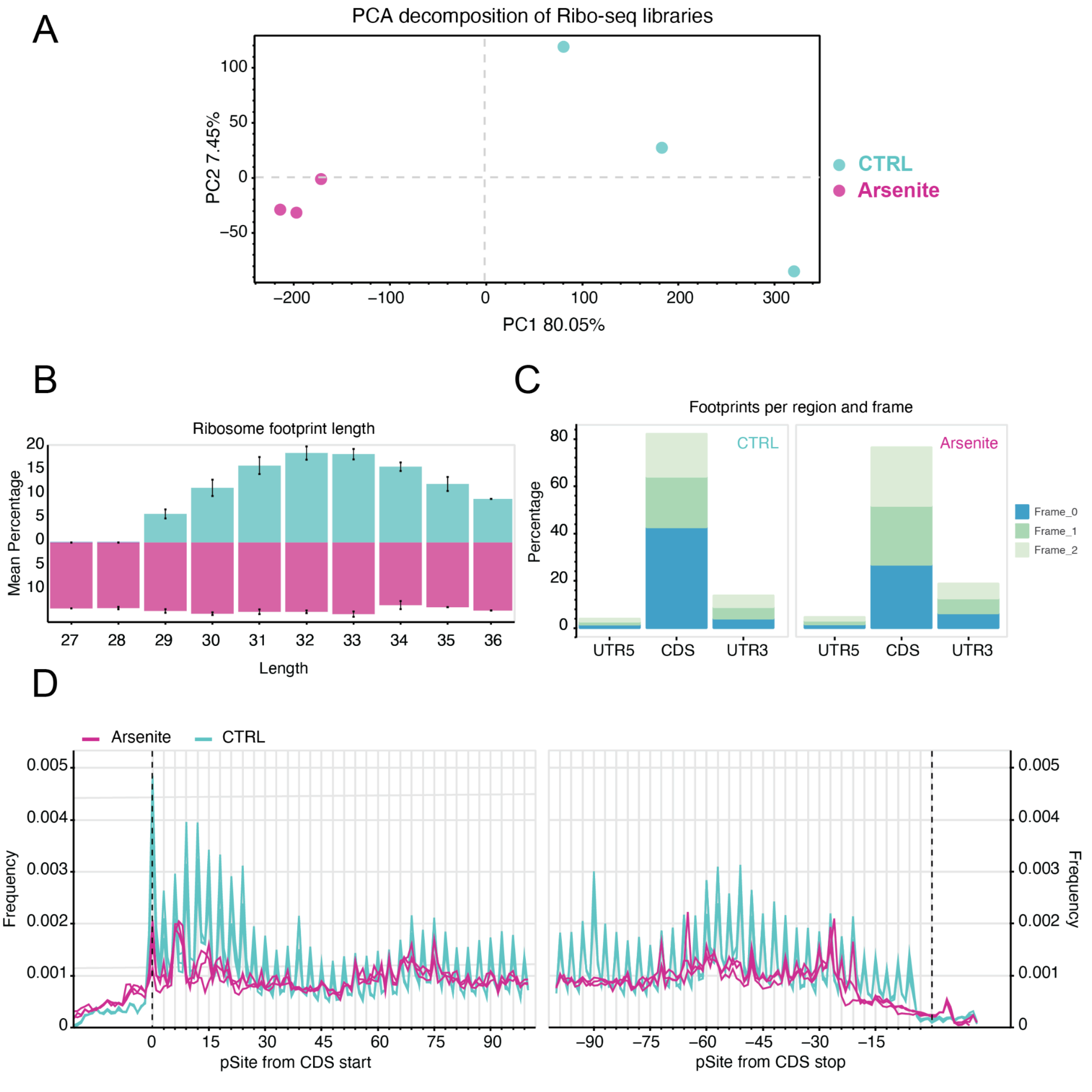

**Figure S17: Modification profiles of full-length tRNAs in arsenite-treated samples show no significant differences by basecalling. (A)** Heatmap of the differential basecalling error of arsenite-treated samples relative to the control, for each nucleotide (*x*-axis) and for each tRNA isoacceptor (*y* axis, ordered in alphabetical order). Upper panel: Ribo-tRNAs, Lower panel: total tRNAs. Each condition was performed with *n* = 3 independent biological replicates.

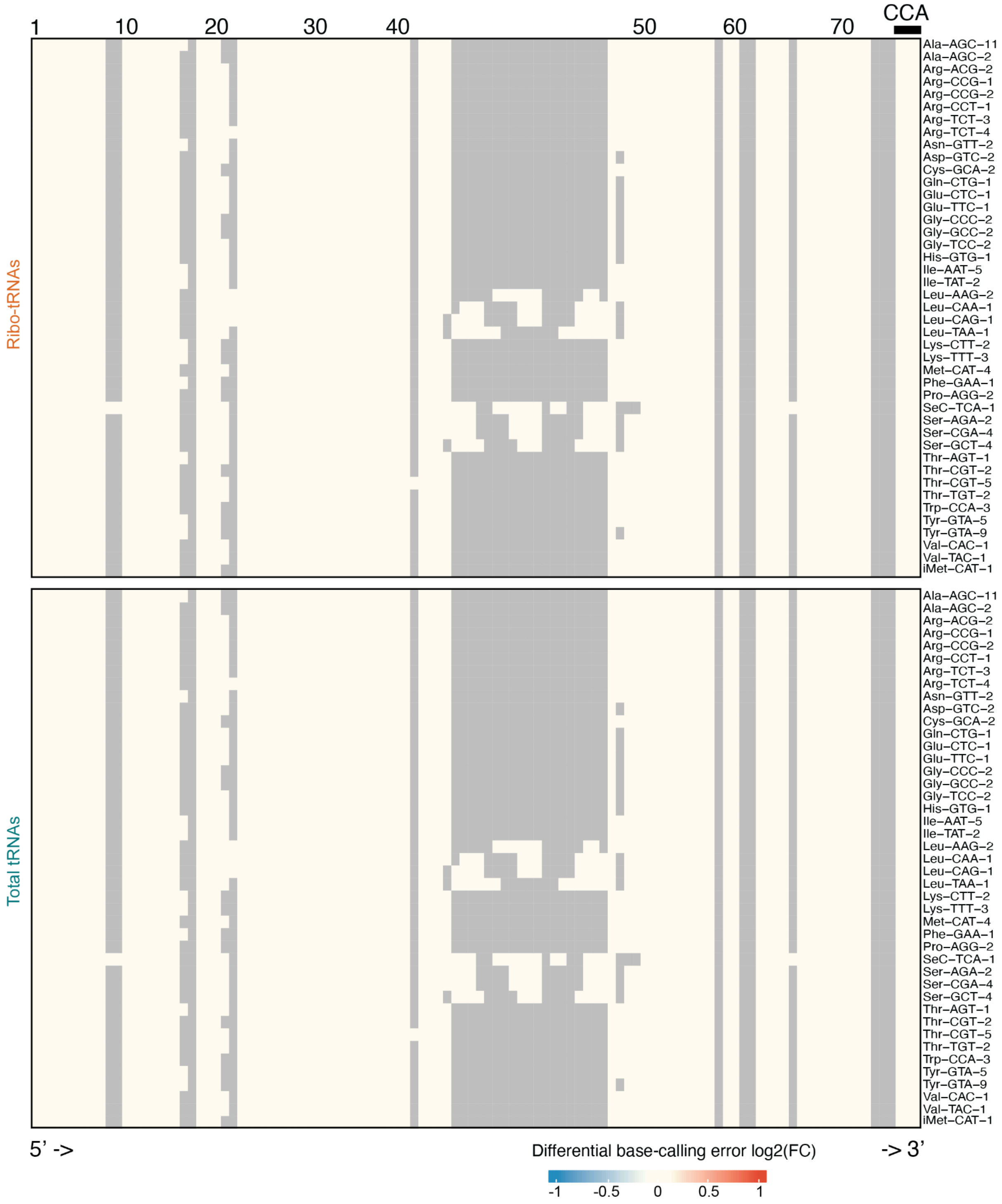

**Figure S18: IGV visualization of significantly fragmented tRNAs.** Integrated Genome Viewer (IGV) tracks displaying tRNA fragmentation profiles upon under control (Ctrl) and arsenite-treated (Ars) conditions, both for ribosome-associated tRNAs (Ribo-tRNA, top) and total tRNAs (Total tRNA, bottom). **(A)** Example IGV snapshots of tRNAs exhibiting a defined fragmentation site upon arsenite exposure. The fragmentation site is indicated by a triangle. Due to nanopore direct RNA sequencing being unable to capture 5’ ends of molecules (it misses the first 12–15 nucleotides), position ~49 the real fragmentation site corresponds to the anticodon region (underlined in black). The positions of the 5′ and 3′ adaptors are shown in green and yellow, respectively. **(B)** Fragmented tRNAs upon arsenite exposure, although without clear identified fragmentation site using the fragmentation analysis pipeline. **(C)** Representative non-fragmented control tRNA (*iMet-CAT*) demonstrating that not all tRNAs undergo fragmentation upon arsenite treatment.

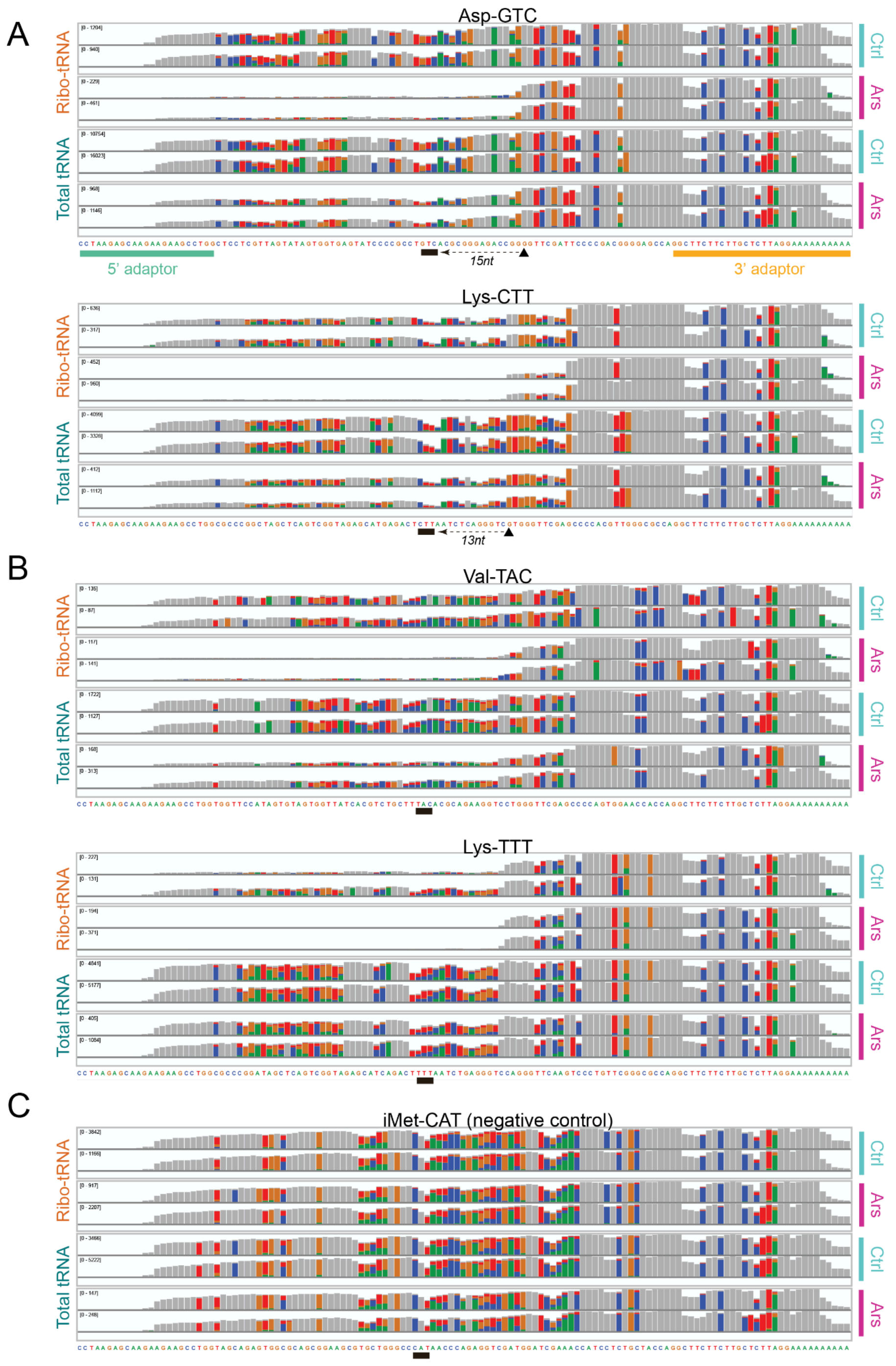

**Figure S19: Arsenite induces specific fragmentation of ribosome-associated tRNAs. (A)** Density plots showing the distribution of read lengths normalized by tRNA length for ribo-tRNAs (top) and total tRNAs (bottom) in control (cyan) and arsenite-treated (pink) samples. Vertical dashed lines mark the mean per condition, with arsenite-treated samples showing shorter read distributions, indicative of increased fragmentation, in both libraries. **(B)** PCA of full-length (FL) and fragmented (FR) tRNA reads across libraries and conditions. Ribo-tRNA profiles (left) show greater separation between control and arsenite-treated samples compared to total tRNA profiles (right), suggesting stronger arsenite-induced fragmentation effects in ribosome-associated tRNAs. (C) *Left:* Schematic outlining the approach for per-site fragmentation analysis. For each nucleotide position within each tRNA, a $2 \times 2$ contingency table was constructed comparing the number of reads terminating versus not terminating at that position in arsenite-treated and control conditions. Statistical testing performed within the edgeR framework (see *Methods*). Right: Volcano plots showing per-site differential fragmentation between arsenite-treated and control samples for ribo-tRNAs and total-tRNAs. Sites with significantly increased fragmentation in arsenite-treated samples are enriched in ribo-tRNAs. Each condition was performed with $n$ = 3 independent biological replicates. For the volcano plot, significance was defined as $p < 0.05$ with an absolute log2 fold-change ≥ 1.

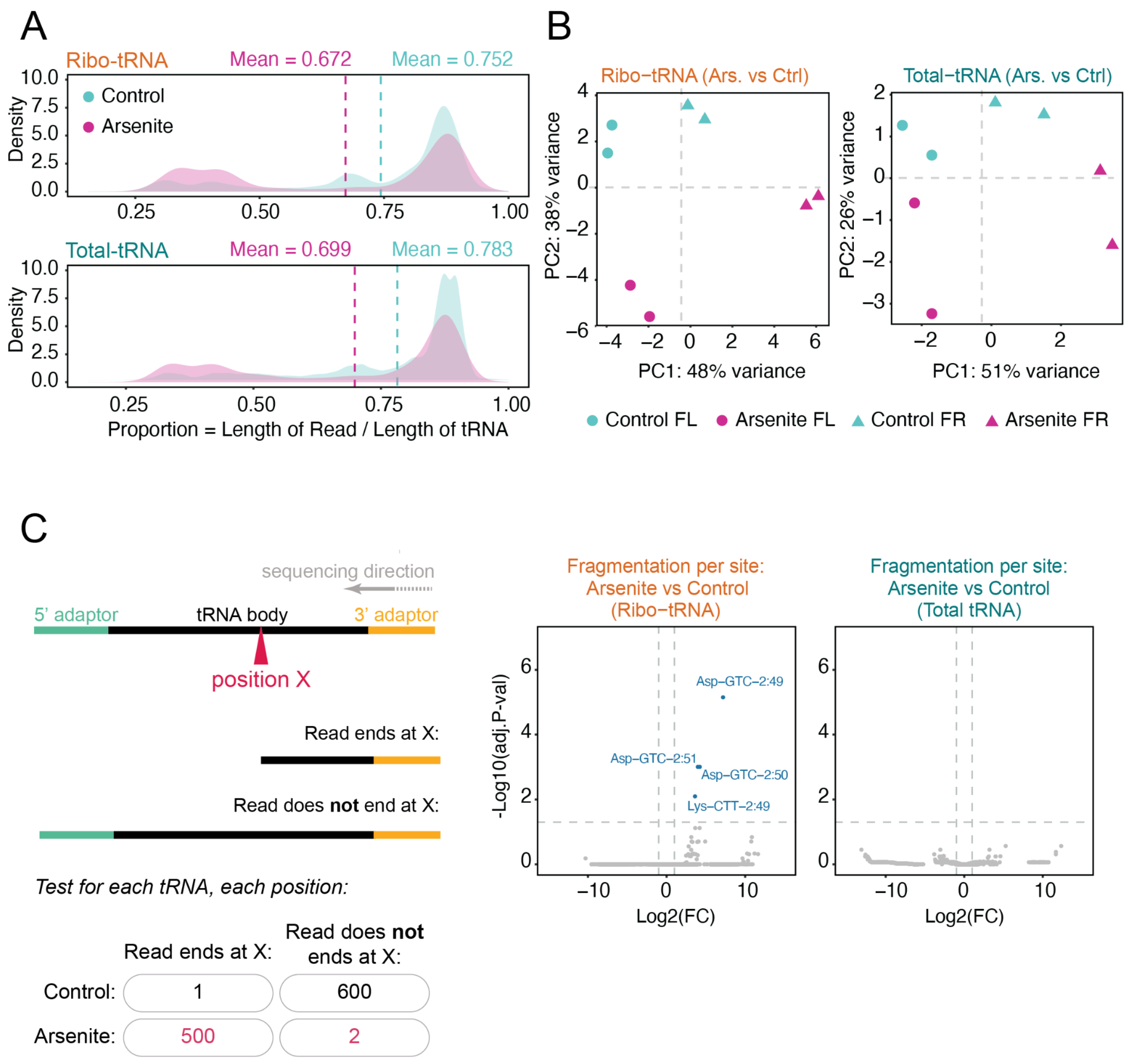

**Figure S20: Fragmentation analysis of additional datasets used in this work.** Fragmentation profiles of ribosome-associated (ribo-tRNA) and total-tRNA populations under various stress conditions, showing that arsenite-induced fragmentation (shown in **Fig. 6F**) is not observed under other perturbations. Harringtonine (A), Arginine starvation (B), Leucine starvation (C) or Met starvation (D). Each condition was performed with *n* = 3 independent biological replicates. Significance is defined as p < 0.05 with an absolute log2 fold-change ≥ 1.

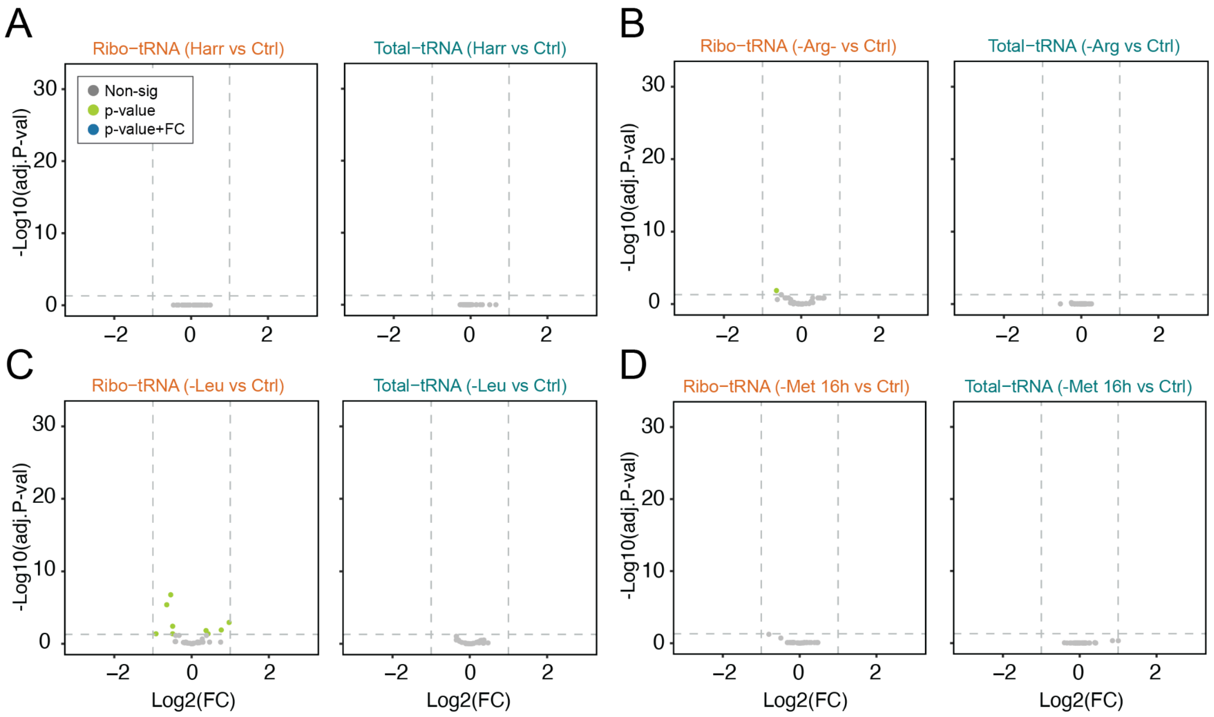

**Figure S21: Cell viability of MCF7 cells is preserved during experimental treatments.** Cells were seeded at $1x10^4$ per well in a 96-well plate and treated as indicated: DMSO, $H_2O$, DMSO + CHX, Ars + CHX, HAR + CHX, or 5% Triton X-100. Sodium arsenite (Ars) was used at 1 mM for 50 minutes, Harringtonine (HAR) at 2 µg/mL for 3 minutes, and CHX at 10 µg/mL for 5 minutes prior to harvest. Cell viability is expressed as the percentage of normalized survival relative to untreated DMEM controls. Error bars represent standard deviation (n = 3). Triton X-100 treatment induced complete cell death (****, P < 0.0001).

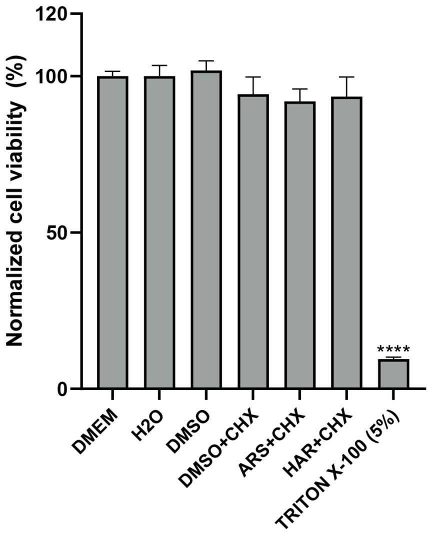
